# Distinct cell-type contributions and network topography of theta-nested gamma oscillations in the medial entorhinal cortex

**DOI:** 10.64898/2026.04.21.719932

**Authors:** Brandon Williams, Riya Dahal, Ananth Vedururu Srinivas, Sheng Xiao, Maria V. Moya, Carmen C. Canavier, Xue Han, Michael N. Economo, John A. White

## Abstract

Theta-nested gamma oscillations in the medial entorhinal cortex (mEC) are essential for spatial coding and memory, but the underlying cellular mechanisms remain unclear. We combined optogenetics, whole-cell electrophysiology, intracellular voltage imaging, and local field potential (LFP) recordings in acute slices from CaMKIIα-ChR2 mice to investigate how excitation and inhibition shape theta-gamma coupling in layer II/III mEC. During theta-frequency stimulation, fast-spiking interneurons received strong gamma-frequency excitation and fired rhythmic bursts, whereas stellate and pyramidal neurons fired more sparsely and were dominated by gamma-frequency inhibition. This sparse firing could support the selective firing of grid cells. Excitatory post-synaptic currents in interneurons preceded inhibitory currents and LFP gamma by ∼3 ms, supporting a pyramidal-interneuron network gamma (PING) mechanism. Pyramidal neurons fired on the descending phase of the gamma cycle, whereas stellate cells and fast-spiking interneurons fired before and after the trough, respectively. Intracellular voltage imaging revealed network gamma synchronization among excitatory neurons at a population level, with topographic clustering of subthreshold membrane potentials, but not spike timing, while individual neurons often skipped gamma cycles. These findings identify the dominant role of reciprocal E-I interactions in generating theta-nested gamma oscillations and highlight distinct cell-type contributions to the temporal dynamics of the mEC. Further, a biophysically realistic computational model predicted gamma cycle skipping in stellate cells and burst firing in fast-spiking interneurons during PING. Our experimental and computational results provide mechanistic insight into how the intrinsic properties of mEC cell types generate oscillatory activity in a manner that could support grid cell function and spatial computation.

**Significance Statement:** Theta-nested gamma oscillations in the medial entorhinal cortex (mEC) are essential for spatial navigation and memory, yet the underlying circuit mechanisms remain incompletely understood. Using optogenetics, voltage imaging, electrophysiology, and biophysically realistic computational modeling, we show that reciprocal interactions between excitatory neurons and fast-spiking interneurons generate robust gamma oscillations through a pyramidal-interneuron network gamma (PING) mechanism. We reveal cell-type specific differences in gamma phase-locking and demonstrate that principal neurons synchronize at the population-level across large laminar distances (up to 800 µm). Further, the voltage activity, but not spike timing, of principal neurons exhibited topographic clustering. These findings clarify how local circuits in mEC generate temporal dynamics critical for grid cell coding and further constrain computational models of spatial representation.

## Introduction

Neural oscillations are critical for coordinating brain activity, enabling information transfer and cognitive function across distributed networks (Buzsáki and Draguhn, 2004; Fries, 2005; Wang, 2010). Cortical gamma oscillations arise from fast inhibitory and excitatory interactions that shape when and which neurons are recruited into local network activity (Bartos et al., 2007; Buzsáki and Wang, 2012). In the medial entorhinal cortex (mEC), gamma (30–140 Hz) rhythms are nested within theta (4–12 Hz) oscillations during navigation and memory-guided behaviors, providing a temporal reference frame for encoding and routing spatial information (Buzsáki and Moser, 2013; Colgin et al., 2009). These theta-nested gamma oscillations are particularly relevant for the temporal coding of grid cells (Brandon et al., 2011; Hafting et al., 2008; Reifenstein et al., 2012).

Two primary models describe how gamma oscillations emerge from cortical microcircuits. In pyramidal-interneuron network gamma (PING), principal neurons excite interneurons, which in turn impose rhythmic inhibitory windows that pace excitatory spiking (Börgers and Kopell, 2003; Traub et al., 1996). In contrast, interneuron network gamma (ING) is driven by mutual inhibition among fast-spiking interneurons, requiring only sustained depolarization (Brunel and Hansel, 2006; Whittington et al., 2000). These mechanisms are not mutually exclusive, and transitions between them may tune oscillatory frequency and interneuron recruitment (Williams et al., 2026).

Principal neurons form local reciprocal connections with fast-spiking interneurons (Buetfering et al., 2014; Couey et al., 2013; Fuchs et al., 2016; Pastoll et al., 2013), which provide fast, rhythmic inhibition critical for maintaining theta-nested gamma oscillations and spatial tuning. Consequently, computational models of grid cell activity and theta-gamma coupling have focused on excitatory-inhibitory circuits (Couey et al., 2013; Pastoll et al., 2013) but often rely on simplified neuron models that neglect the intrinsic dynamics of individual cell types. As a result, it remains unclear how the intrinsic properties of different excitatory neurons shape PING dynamics, and how these cell-type-specific interactions give rise to the local field potential (LFP) signal that reflects network gamma oscillations. Establishing this link between synaptic timing, cell-type recruitment, and the LFP is essential for interpreting population-level measurements of theta-nested gamma activity.

Although gamma generation is increasingly understood, less is known about how distinct excitatory cell types participate. Principal neurons in layer II/III mEC consist of stellate and pyramidal neurons (Alonso and Klink, 1993; Canto and Witter, 2012), which differ in intrinsic conductances and dendritic integration (Haas et al., 2007; Haas and White, 2002), while both contribute to grid cell coding (Domnisoru et al., 2013). These differences may influence how inhibitory synchronization shapes excitatory output, yet direct measurements of their gamma-locked recruitment have been limited. Previously, individual stellate cells and fast-spiking interneurons have been shown to exhibit relatively strong gamma phase locking, but this is diminished at the population level, contrasting with model predictions (Pastoll et al., 2013). The gamma phase-locking properties of pyramidal cells remain unexplored, and improved models are necessary to be consistent with experimental recordings.

Excitatory neurons in the mEC are spatially organized, with stellate cells interspersed between patches of pyramidal cells (Kitamura et al., 2014; Ray et al., 2014). Grid cells also exhibit anatomical clustering (Gu et al., 2018). These ensembles reside within a dense mesh of PV+ interneurons whose synaptic and electrical coupling decreases sharply with distance (Fernandez et al., 2022; Huang et al., 2024), consistent with local modules of excitatory-inhibitory connectivity (Huang et al., 2024). This arrangement implies that gamma-frequency inhibition may not be distributed uniformly across the local network but instead may selectively engage subsets of nearby excitatory neurons. However, the extent to which gamma-synchronous activity is spatially structured across excitatory populations remains unclear.

Here, we used optogenetic theta drive, whole-cell patch clamp, LFP recordings, and intracellular voltage imaging in acute slices from CaMKIIα-ChR2 mice to investigate theta-nested gamma dynamics in layer II/III mEC. We show that synaptic and spike timing support a PING mechanism, and that distinct excitatory cell types exhibit differential participation in gamma oscillations. Furthermore, population voltage imaging reveals gamma-synchronous activity across large spatial scales with modular clustering of subthreshold dynamics. Together, these findings provide a unified view of how cell-type-specific interactions and spatial organization shape theta-nested gamma oscillations in the mEC.

## Results

Here we examine an *in vitro* analogue of theta nested gamma in which gamma oscillations are an emergent property of the mEC network when the theta drive is directed primarily to the excitatory cells. CaMKIIα is mainly expressed in principal neurons in the parahippocampal region, albeit with measurable crossover into PV+ cells in regions CA1 and CA3 (Butler et al., 2018). A transgenic CaMKIIα-Cre mouse line was crossbred with a transgenic LoxP-ChR2-LoxP-EYFP mouse line to constitutively express ChR2-EYFP in CaMKIIα+ neurons (CaMKIIα-ChR2 mice). Like (Butler et al., 2018), we found dense expression of ChR2-EYFP in layer II/III mEC. Acute brain slices were prepared from adult male and female CaMKIIα-ChR2 mice in roughly equal numbers. Whole cell patch clamp recordings were obtained from stellate, pyramidal and fast-spiking interneurons in layer II/III mEC during 8 Hz sinusoidal optogenetic stimulation, to simulate network theta drive observed in spatial behaviors.

### Firing properties of stellate, pyramidal and fast-spiking interneurons during theta-frequency optogenetic drive of CaMKIIα+ neurons

Intracellular voltage recordings captured the responses of mEC neurons during theta frequency optogenetic CaMKIIα+ stimulation (Fig. 1A). Cell types, determined using established electrophysiological signatures (Materials and methods), showed repeatable responses (Fig. 1B). Per theta cycle, stellate cells fired a median of 1.15 spikes (IQR: 0.24–1.89, n = 16), pyramidal cells fired 1.10 spikes (1–2.29, n = 24), and fast-spiking interneurons fired 2.43 spikes (1.03–3.34, n = 9). Interestingly, stellate and pyramidal cells appeared to fire more spikes before peak theta stimulation, while fast-spiking interneurons sustained firing after peak stimulation (Fig. 1C). Stellate and pyramidal cells mostly fired at rates below 70 Hz, lower than gamma-frequency inputs seen in intracellular recordings (70–110 Hz; see Fig. 2 below). This result suggests the possibility of gamma cycle skipping (Fig. 1D). In contrast, fast-spiking interneurons fired bursts at rates up to 300 Hz, indicating multiple spikes per putative gamma cycle. Together, these results suggest that excitatory neurons maintain network gamma oscillations by alternating subpopulations on different gamma cycles. Fast-spiking interneurons provide fast feedback inhibition, sometimes firing bursts, to tightly control excitation in mEC through a PING mechanism.

**Figure 1:**
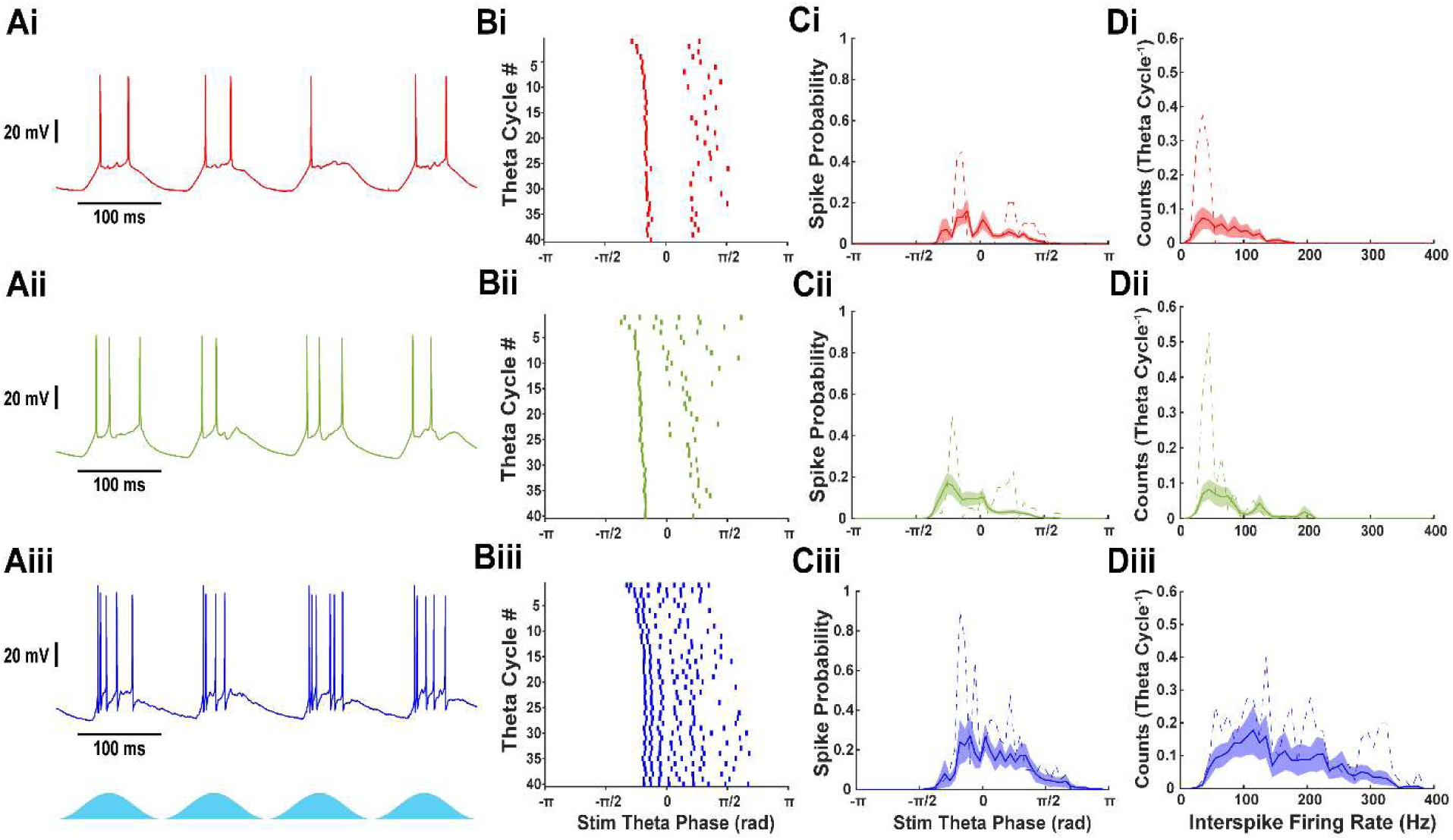
Fast-spiking interneurons fire in gamma frequency bursts, while stellate and pyramidal cells fire more sparsely. A) Voltage traces from (i) stellate, (ii) pyramidal, and (iii) fast-spiking interneurons during optogenetic stimulation of CaMKIIα+ neurons. B) Raster plots showing 40 theta periods of sinusoidal 8 Hz optogenetic stimulation for the same recording as A. C) Spike probability histogram for each neuron relative to phase of theta stimulation [mean (solid line) ± S.E.M. (shaded region)]. Example spike probability from neurons shown in A and B (dashed line). D) Interspike firing rate histogram for each neuron [mean (solid line) ± S.E.M. (shaded region)]. Example spike probability from neurons shown in A and B (dashed line).

**Figure 2:**
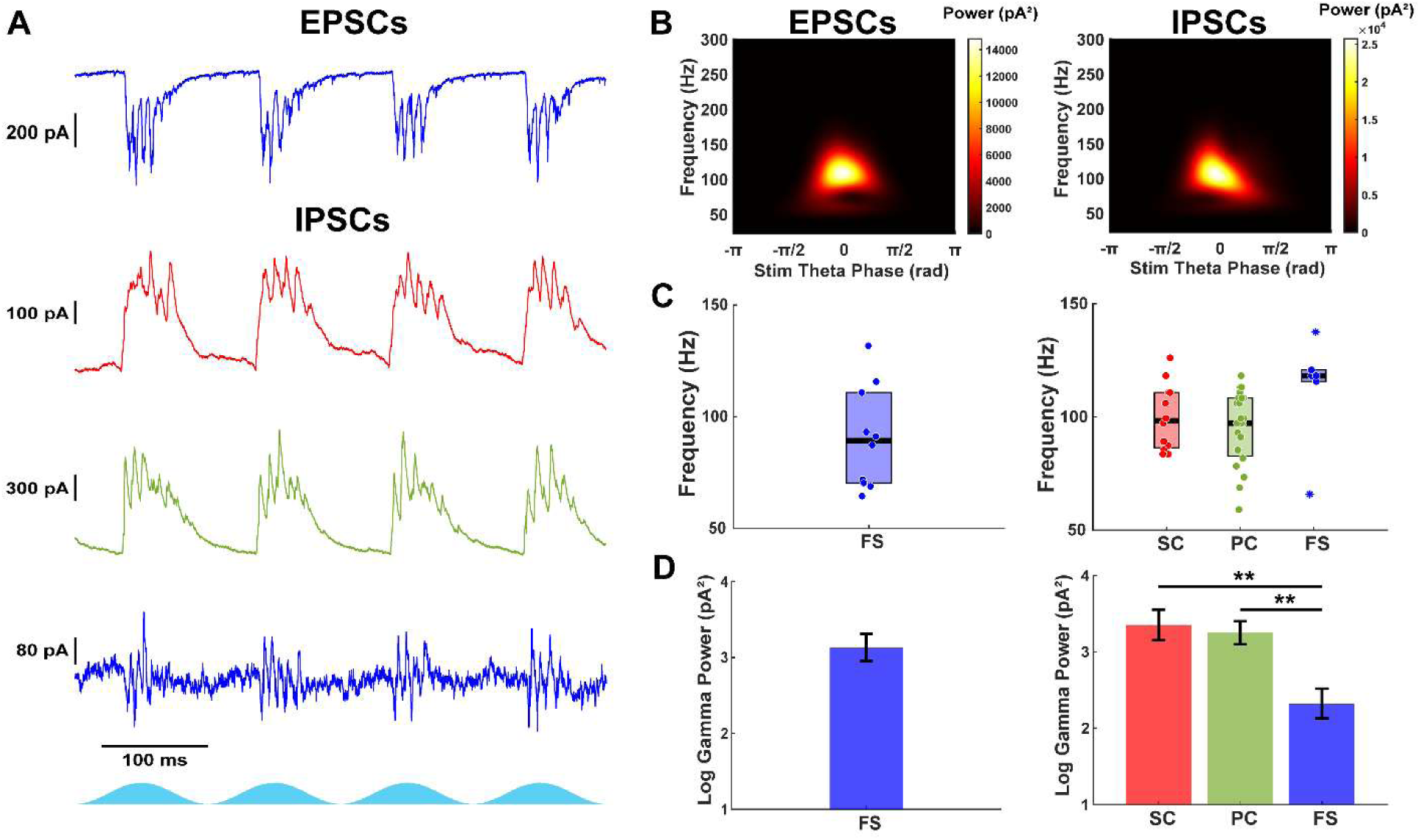
Fast-spiking interneurons receive strong excitation and moderate inhibition, while stellate and pyramidal cells receive strong inhibition. A) Example post-synaptic currents recorded during optogenetic stimulation of CaMKIIα+ neurons. From top to bottom: EPSCs from fast-spiking interneuron. IPSCs recorded from stellate, pyramidal, and fast-spiking interneuron. B) Average scalogram of gamma-filtered PSCs from neurons in A. Left: EPSCs from fast-spiking interneuron. Right: IPSCs from pyramidal cell. C) Gamma frequency of PSCs from all neurons. Left: EPSCs from fast-spiking interneurons. Right: IPSCs from stellate, pyramidal, and fast-spiking interneurons. D) Peak gamma power of PSCs from all neurons. Left: EPSCs from fast-spiking interneurons. Right: IPSCs from stellate, pyramidal, and fast-spiking interneurons.

### Theta-nested gamma oscillations are observed in excitatory and inhibitory post-synaptic currents in layer II/III mEC during CaMKIIα+ stimulation

Previously, (Butler et al., 2018) demonstrated that optogenetically driven theta-nested gamma oscillations in the mEC of CaMKIIα-ChR2 mouse brain slices were reduced during the blockade of AMPA/kainate receptors, suggesting that fast excitatory synaptic currents can significantly strengthen gamma oscillations. In contrast, the blockade of NMDA receptors did not change the power of theta-nested gamma during theta frequency optogenetic drive, suggesting that the primary excitatory contribution to the gamma oscillations is mediated by AMPA/kainite receptors. To investigate the differential contributions of excitation versus inhibition in individual cell types, we attempted to isolate excitatory and inhibitory membrane currents in mEC neurons by voltage clamping at −70 mV and 0 mV, respectively.

After log-transformation, cell type was found to significantly influence the peak gamma power of currents recorded at −70 mV (F_(2,40)_ = 25.04, p = 8.88e−8, one-way ANOVA). Fast-spiking interneurons displayed strong gamma-frequency EPSCs (geometric mean peak gamma power: 1362 pA², 95% CI: 515–3599, n = 10; Fig. 2A, B, D). In contrast, stellate cells (108 pA², 95% CI: 59–201, n = 12, p = 5.94e−5, Tukey-Kramer) and pyramidal cells (50 pA², 95% CI: 28–90, n = 21, p = 4.86e−8) showed significantly weaker EPSCs, consistent with prior work indicating minimal recurrent excitation in superficial mEC (Couey et al., 2013; Fuchs et al., 2016). The frequency of EPSCs in fast-spiking interneurons was in the fast gamma range (median: 89.2 Hz, IQR: 70.3–110.8, n = 10/10; Fig. 2C).

Gamma-frequency IPSCs were present in all cell types (Fig. 2A–D). Cell type was found to significantly affect IPSC peak gamma power (F_(2,41)_ = 6.55, p = 0.0034, one-way ANOVA). Fast-spiking interneurons exhibited weaker IPSCs (geometric mean: 210 pA², 95% CI: 70–631, n = 9; Fig. 2D) compared to stellate (2251 pA², 95% CI: 773–6556, n = 12, p = 0.0062) and pyramidal cells (1775 pA², 95% CI: 850–3709, n = 23, p = 0.0057). IPSCs did not differ between stellate and pyramidal cells (p = 0.91). The frequency of IPSCs was not statistically different across groups (stellate: 98.3 Hz, IQR: 86.3–110.8, n = 12/12; pyramidal: 97.3 Hz, IQR: 82.7–108.4, n = 23/23; fast-spiking: 118.2 Hz, IQR: 115–120, n = 6/9; χ²_(2,38)_ = 5.81, p = 0.055, Kruskal-Wallis). Recordings with low gamma power, low signal-to-noise, and high spectral bandwidth were removed from frequency analysis (see Materials and methods).

The similarity between IPSC frequencies in excitatory cells and EPSC frequencies in fast-spiking interneurons implies that they are participating in the same pattern of rhythmic activity. Thus, we conclude that a PING circuit mechanism dominates during optogenetic activation of local excitatory neurons in the mEC.

### Excitation precedes inhibition and the local field potential oscillation

Theta-nested gamma oscillations have been observed in layer II/III mEC during various behaviors. Previously, optogenetic stimulation of Thy1+ neurons in layer II mEC generated theta-nested gamma oscillations in the LFP, IPSCs in stellate cells, and EPSCs in fast-spiking interneurons (Pastoll et al., 2013). Thy1 expression is primarily found in stellate cells and fast-spiking interneurons. Another study reported theta-nested gamma oscillations in the LFP during optogenetic stimulation of CaMKIIα+ neurons (Butler et al., 2018), which include both stellate and pyramidal cells—two principal cell types that exhibit grid cell tuning (Domnisoru et al., 2013). Notably, in this context, inhibitory interneurons are recruited indirectly via excitation, consistent with a PING (pyramidal-interneuron network gamma) mechanism. However, that study did not examine the activity of individual cell types (Butler et al., 2018). Building on this work, we optogenetically stimulated CaMKIIα-expressing neurons to determine the strength and timing of synaptic inputs, and spike timing of individual cell types in mEC during PING. Further, we aim to determine if gamma oscillations generated through indirect activation of local interneurons (CaMKIIα+) are different from direct stimulation of excitatory and inhibitory neurons (Thy1+).

A local field potential electrode was inserted into layer II/III mEC to record network theta-nested gamma oscillations. During simultaneous whole-cell voltage clamp, LFP recording and optogenetic stimulation of CaMKIIα+ neurons, excitatory and inhibitory membrane currents were measured across different cell types (Fig. 3A). Gamma-frequency activity was evident in both the membrane currents of mEC neurons and the LFP (Fig. 3B, C). Cross-correlation coefficient analysis revealed strong alignment between IPSCs and the LFP across all cell types: stellate cells (median: 0.77, IQR: 0.77–0.79, n = 6), pyramidal cells (median: 0.62, IQR: 0.53–0.76, n = 18), and fast-spiking interneurons (median: 0.56, IQR: 0.45–0.72, n = 5; Fig. 3F). No significant differences were found between the correlation coefficients in each group (stellate vs. pyramidal vs. fast-spiking: χ^2^_(2,26)_ = 3.57, p = 0.17; Kruskal-Wallis test). Similarly, EPSCs in fast-spiking interneurons were strongly correlated with the LFP (median: 0.58, IQR: 0.48–0.68, n = 8; Fig. 3E). EPSCs in excitatory neurons were not analyzed because they were too small to be accurately measured.

**Figure 3:**
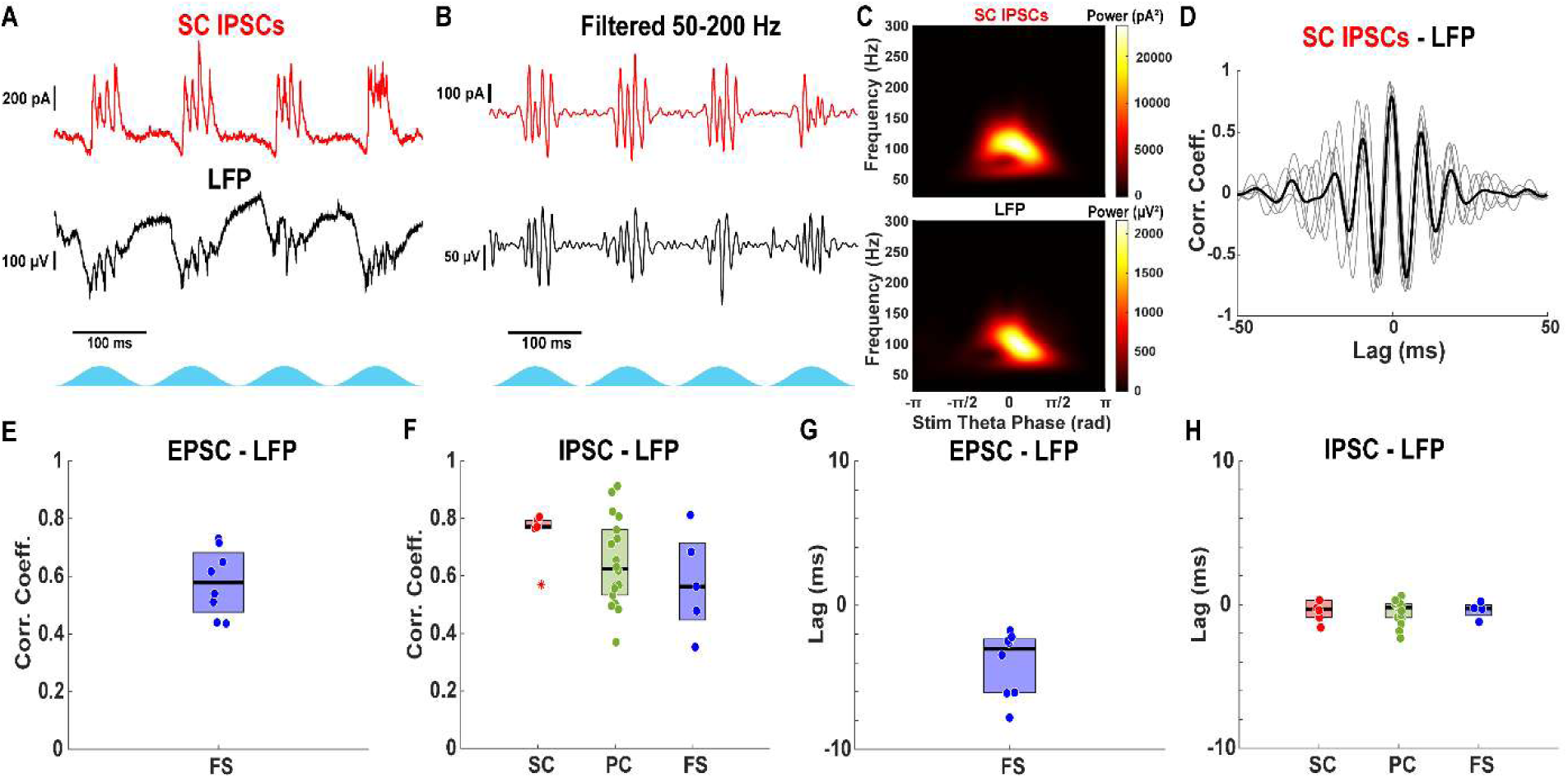
EPSCs precede IPSCs and LFP during theta frequency optogenetic stimulation of CaMKIIα+ neurons. A) Example IPSCs received by stellate cell (red) and concurrent LFP (black). B) Gamma frequency filtered IPSCs (red) and LFP (black) from A. C) Average scalogram of gamma filtered IPSCs (red) and LFP (black) from B demonstrating similar gamma frequency oscillations. D) Average cross correlogram (black) between gamma filtered currents and LFP from B and individual theta cycles (gray). IPSCs are highly correlated with the LFP with no lag, demonstrating that the LFP oscillation is likely driven by local inhibitory currents. E) Average peak cross correlation coefficient between EPSCs and concurrent LFP. EPSCs received by fast-spiking interneurons are highly correlated with the LFP, indicating a strong PING mechanism. F) Average peak cross correlation coefficient between IPSCs received by stellate (red), pyramidal (green) and fast-spiking interneurons (blue) and the concurrent LFP. IPSCs are highly correlated with the LFP. G) Average peak correlation lag between EPSCs and LFP. The LFP lags the EPSCs by approximately 3 ms. H) Average peak correlation lag between IPSCs and LFP. The LFP is oscillating approximately at the same time as the IPSCs.

The peak correlation between IPSCs and the LFP occurred with negligible temporal lag across all cell types (stellate: median: –0.33 (IQR: −0.90–0.30) ms, n = 6; pyramidal: –0.20 (−0.91–0.05) ms, n = 17; fast-spiking: –0.28 (−0.75–−0.03) ms, n = 4; Fig. 3H). No significant differences were found between the correlation lags in each group (stellate vs. pyramidal vs. fast-spiking: F_(2,24)_ = 0.03, p = 0.97; one-way ANOVA test). In contrast, EPSCs in fast-spiking interneurons preceded the LFP by ∼3 ms (median: −3.03 (−6.08–−2.35) ms, n = 8; Fig. 3G). Consistent with a PING mechanism in which phasic input from the excitatory cells evokes a population burst in the inhibitory cells, we show that fast-spiking interneurons receive excitatory input approximately 3 ms before inhibitory feedback arrives during each gamma cycle. Overall, the LFP closely reflects the timing and frequency of local inhibitory currents in the mEC.

### Cell-type specific temporal organization of gamma phase locking in mEC

Theta-nested gamma oscillations in the mEC are thought to reflect coordinated excitation-inhibition dynamics, yet the relative timing of different cell types within the gamma cycle remains unresolved. To quantify cell-type specific phase locking to gamma oscillations, we performed simultaneous intracellular voltage recordings and LFP measurements in stellate cells, pyramidal cells, and fast-spiking interneurons during theta-frequency CaMKIIα optogenetic stimulation (Fig. 4A).

**Figure 4:**
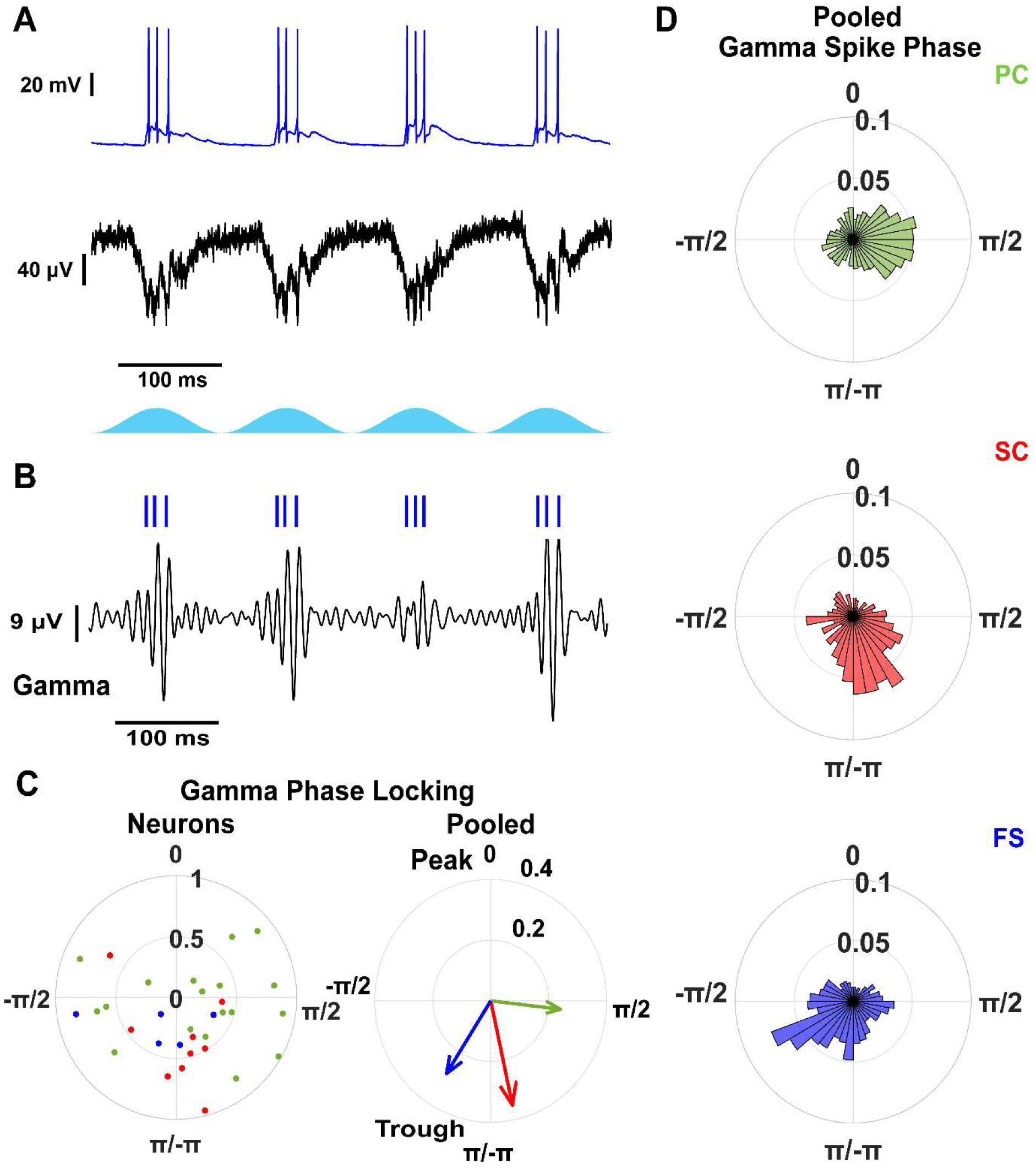
mEC neurons are phase locked to gamma oscillations. A) Example voltage trace of a fast-spiking interneuron (blue) and concurrent LFP (black) during optogenetic stimulation of CaMKIIα+ neurons. B) Filtered gamma frequency LFP component (black) and raster plot of fast-spiking interneuron (blue) from A. C) Resultant vector comparison between stellate (red), pyramidal (green), and fast-spiking interneurons (blue). Left: Individual neurons. Right: Pooled within cell types. D) Pooled gamma spike phase distributions for stellate, pyramidal and fast-spiking interneurons.

At the single-cell level, vector strength did not differ across cell types (stellate: 0.55 (0.44–0.67), n = 9; pyramidal: 0.58 (0.47–0.70), n = 18; fast-spiking: 0.43 (0.24–0.62), n = 5; p = 0.47, one-way ANOVA; Fig. 4B), indicating comparable phase locking across neurons. However, preferred phase distributions differed. Most pyramidal neurons (13/18) fired on the descending phase, with a smaller subgroup (5/18) on the ascending phase, whereas stellate cells and fast-spiking interneurons exhibited unimodal phase preferences near the trough.

When pooling spikes within each cell type, stellate cells exhibited higher vector strength than fast-spiking interneurons and pyramidal cells (stellate: 0.35; fast-spiking: 0.28; pyramidal: 0.24; Fig. 4B–D). Pyramidal cell spikes occurred earlier in the gamma cycle (1.69 (1.50–1.87) rad) than stellate cells (2.93 (2.77–3.08) rad) and fast-spiking interneurons (−2.57 (−2.79–−2.36) rad). The phase difference between pyramidal cells and fast-spiking interneurons (∼3.2 ms for a 10 ms gamma cycle) matched the delay between excitation and inhibition measured in fast-spiking interneurons relative to the LFP (Fig. 3).

To assess phase locking across spikes within theta cycles, we used pairwise phase consistency (PPC; (Vinck et al., 2010)), which is robust to differences in spike count. All cell types showed strong theta phase locking across spikes (Fig. S1A). In contrast, gamma phase locking declined across successive spikes in stellate and pyramidal cells but remained stable in fast-spiking interneurons (stellate: χ^2^_(2,17)_ = 14.37, p = 7.57e–4; pyramidal: χ^2^_(2,31)_ = 8.70, p = 0.013; fast-spiking: χ^2^_(3,13)_ = 1.03, p = 0.79; Kruskal-Wallis test; Fig. S1B). Stellate cells showed a rapid reduction after the first spike, pyramidal cells after the second spike, whereas fast-spiking interneurons maintained consistent gamma phase locking across spikes.

Overall, these results suggest cell-type specific temporal organization within the gamma cycle, with fast-spiking interneurons maintaining consistent gamma coupling and excitatory neurons exhibiting spike-dependent reductions in gamma phase locking.

### Voltage imaging reveals network synchronization at gamma frequencies with no linear distance dependence between spike correlations across layer II/III mEC

Genetically encoded voltage indicators (GEVIs) offer high temporal resolution and cell-type specificity for monitoring membrane potential dynamics. Recent advances in GEVIs and optical techniques now allow for the simultaneous imaging of dozens of neurons during optogenetic stimulation (Abdelfattah et al., 2023; Xiao et al., 2024). Unlike traditional electrophysiology, intracellular voltage imaging preserves 2D spatial information, enabling the cell-type specific study of anatomical organization in neural activity. While whole-cell recordings provide precise measurements of synaptic timing in individual neurons, they do not capture how activity is coordinated across populations. Here, we used large field-of-view voltage imaging to determine whether gamma synchronization emerges locally or across distributed excitatory networks, and whether this activity exhibits spatial organization.

As an alternative to simply recording repeated trials in single neurons, we investigated the spatial and network dynamics underlying theta-nested gamma oscillations in mEC by simultaneously imaging the intracellular voltage of dozens of layer II/III neurons during theta frequency optogenetic stimulation of CaMKIIα+ neurons and LFP recording (Fig. 5A, B). Post-hoc immunohistology revealed minimal expression of our voltage sensor, Voltron2, in PV+ interneurons (Fig. S2), limiting our analysis to putative excitatory neurons. We hypothesize that the tropism of the AAV2retro virus restricts transfection of PV+ interneurons in mEC.

**Figure 5:**
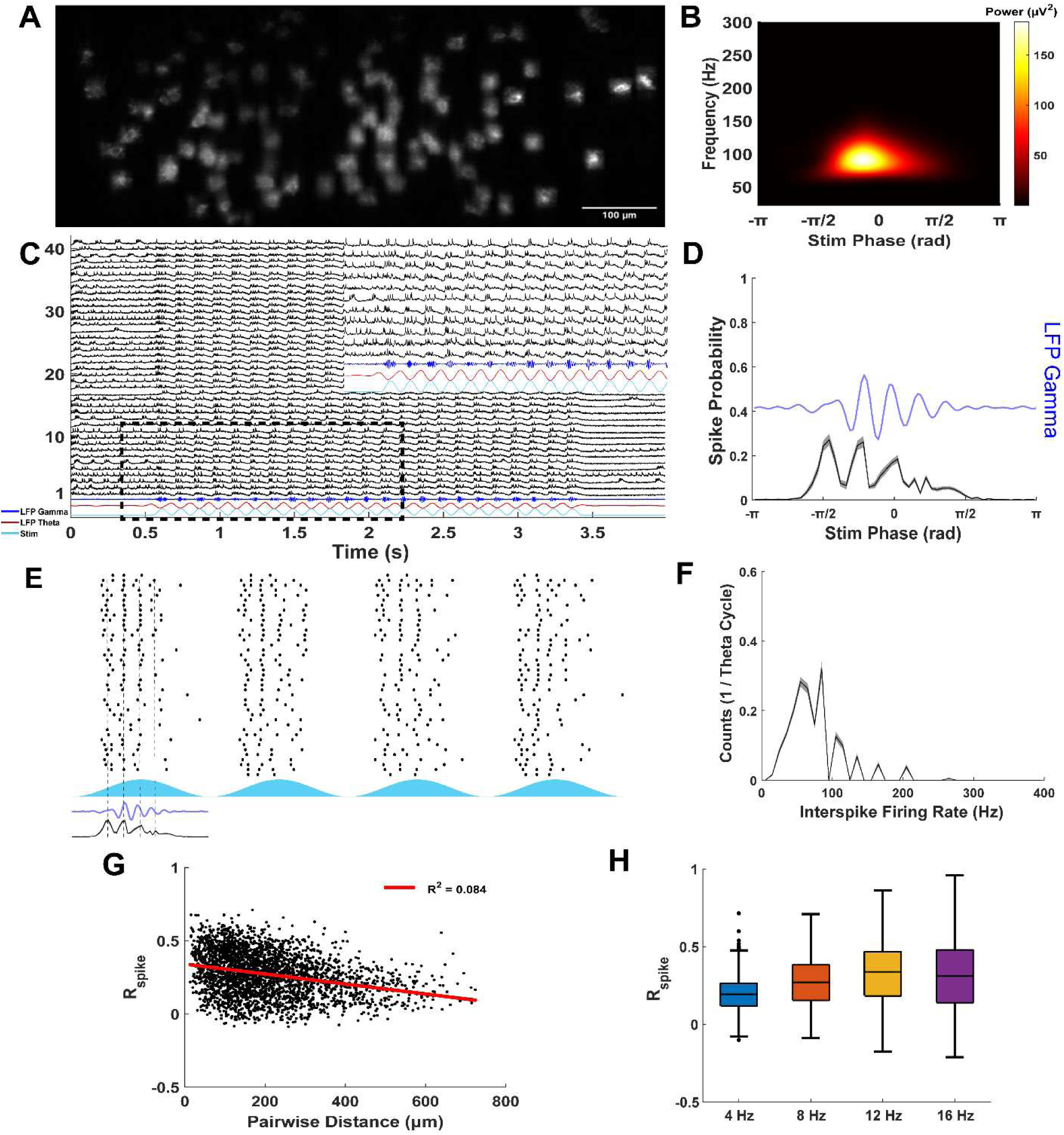
Voltage imaging reveals network dynamics of layer II/III mEC neurons. A) Example field of view for targeted illumination voltage imaging. Fluorescent neurons express Voltron2. B) Average LFP Scalogram demonstrates strong gamma frequency activity. C) Voltage traces from 41 neurons shown in A and the simultaneous LFP activity. Inset zooms in on traces within dashed rectangle. D) Average spike timing of all neurons in C (n = 41) relative to theta stimulation phase [mean (black) ± S.E.M. (gray)]. Average LFP gamma (blue) lags excitatory neuron spiking oscillations. E) Spike raster plot (black) of all neurons in C (n = 41) during 4 consecutive cycles of 8 Hz optogenetic stimulation. Light stimulus is shown in light blue. Average LFP (blue) and spike histogram [mean (black) ± S.E.M. (gray)] aligned with example theta period using dashed lines. F) Average interspike firing rate histograms per theta stimulation period from neurons in all imaging sessions (n = 240). G) Relationship between spike train correlations of all neurons (n = 240) and pairwise spatial distance. H) Comparison of spike train correlations for all neurons (n = 240) across different theta stimulation frequencies.

To determine whether nested gamma frequency was an emergent network property independent of the frequency of theta drive, as expected with a PING mechanism, a local field potential electrode was positioned in layer II/III to capture network-level theta-nested gamma oscillations (Fig. 5B, C). We also varied the frequency of optogenetic stimulation (4, 8, 12, and 16 Hz) to assess frequency-dependent network dynamics. As stimulation frequency increased, the number of gamma cycles per theta period decreased, while the gamma frequency remained constant (Peak Gamma Frequency during 4, 8, 12, and 16 Hz stim: χ^2^_(3,39)_ = 3.71, p = 0.29; Fig. S3), indicating that gamma frequency is governed by local E-I interactions (i.e., PING) rather than the external drive. Consistent with this interpretation, excitatory neurons exhibited synchronized activity at gamma frequencies across a large field of view (up to ∼800 µm along the medial-lateral axis; Fig. 5D, E), demonstrating that gamma coordination is widely distributed across the local network rather than confined to small spatial domains. Gamma frequency oscillations in the LFP followed excitatory firing and likely reflect local inhibition, as demonstrated in Fig. 3.

Across all imaging sessions, excitatory neurons fired at higher rates (8 Hz stim: median: 2.76, IQR: 2.02–3.38 spikes per stim period, n = 240) than in whole-cell recordings, likely due to the additional ChR2 current induced by imaging with a high power 561 nm laser. However, as in whole-cell recordings (Fig. 1), individual neurons fired at fast gamma frequencies or slower (Fig. 5F), consistent with gamma cycle skipping at the single-cell level despite population-level synchronization.

Although spike timing was strongly phase locked to the LFP theta rhythm, we did not find gamma phase locking among excitatory neurons at 8 Hz theta frequency drive (Fig. S4). It’s possible that fast gamma phase locking could not be resolved at the 800 Hz imaging rate. In addition, the lower SNR of voltage imaging and filtering of voltage activity due to the kinetics of Voltron2 could contribute to the variability in spike timing compared to whole-cell recordings.

Interestingly, we found a trend of stronger gamma phase locking for the first spike in each theta period when increasing the frequency of sinusoidal optogenetic stimulation to 12 or 16 Hz (Fig. S5), suggesting that faster activation of excitatory neurons reduces spike timing variability enough to resolve gamma phase locking at an 800 Hz frame rate.

Intracellular voltage imaging advantageously permits the analysis of spatial dynamics across layer II/III mEC. Pairwise spike correlations showed no linear dependence on the distance between neuron pairs (Linear Regression R^2^: 4 Hz: 0.071, n = 2821; 8 Hz: 0.085, n = 2821; 12 Hz: 0.060, n = 1408; 16 Hz: 0.069, n = 1491; Fig. 5G), indicating that temporal spike patterns are not spatially clustered. Furthermore, pairwise spike correlations increased with stimulation frequency (Spike Correlation R_spike_: 4 Hz: 0.19 (0.12–0.26), n = 2821; 8 Hz: 0.27 (0.16–0.38), n = 2821; 12 Hz: 0.34 (0.18–0.46), n = 1408; 16 Hz: 0.31 (0.14–0.48), n = 1491; Fig. 5H), likely reflecting enhanced firing synchrony due to sharper temporal input and a reduced total number of spikes.

Together, these results demonstrate that gamma oscillations in mEC are coordinated across distributed excitatory populations without spatial clustering of spike patterns. This suggests that local circuit interactions can support large-scale temporal coordination, while the precise recruitment of individual neurons remains heterogeneous.

### Subthreshold activity is correlated among clusters of excitatory neurons across layer II/III mEC

The organization of spatial firing properties in mEC neurons is an important constraint for developing models of grid cells. Networks of grid cells form anatomically overlapping, but functionally discrete modules that share grid spacing and orientation but have distributed spatial phases to represent the entire environment (Barry et al., 2007; Hafting et al., 2005; Shipston-Sharman et al., 2016; Stensola et al., 2012). In layer II, pyramidal (calbindin+) cells have shown to form compact anatomical clusters (Island cells’) while stellate (reelin+) cells are more sparsely distributed between these clusters [Ocean cells; (Kitamura et al., 2015; Ray et al., 2014; Varga et al., 2010)]. Grid cells also display physical clustering and are functionally organized by spatial firing phase (Heys et al., 2014), although this anatomical organization is weak compared to the neural sheets in continuous attractor models (Burak and Fiete, 2009; Couey et al., 2013; Fuhs and Touretzky, 2006; Guanella et al., 2007; Pastoll et al., 2013). Here, we asked whether neuronal activity during theta-nested gamma oscillations is anatomically organized across excitatory populations, and whether this organization differs between subthreshold membrane dynamics and spike output.

Using the example recording in Fig. 5, a correlation matrix was computed to examine the similarity of voltage signals between mEC neurons during theta-frequency stimulation (Fig. 6A). Inspection of the correlation heatmap revealed a subset of neurons that were weakly correlated with the rest of the population, possibly reflecting cells outside the CaMKIIα+ excitatory population. In contrast, a large group of neurons exhibited strong correlations in their voltage activity. Because neurons were ordered along the medial-lateral axis, this pattern suggested that functionally correlated neurons may be spatially clustered.

**Figure 6:**
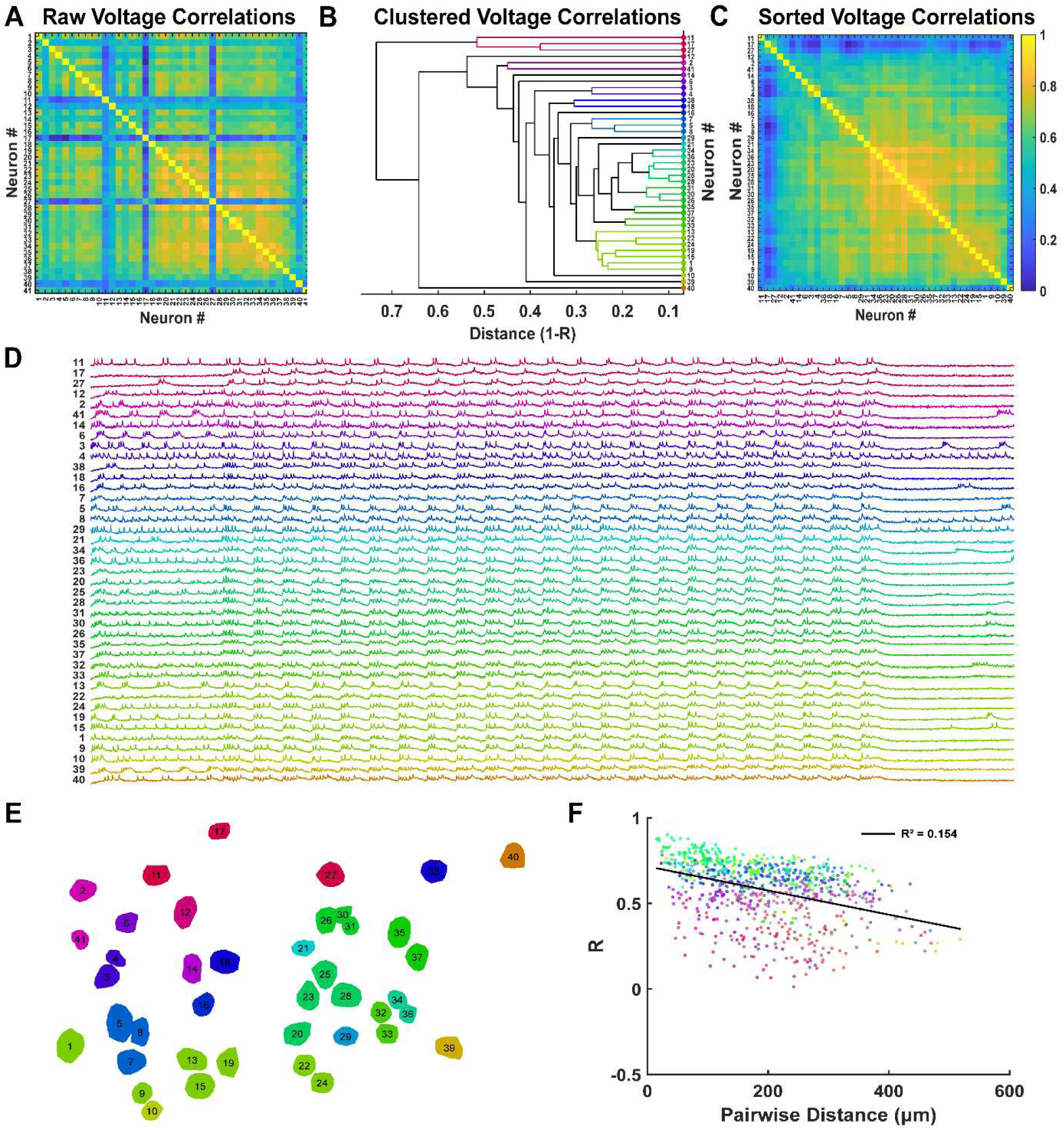
Local clusters of excitatory mEC neurons exhibit correlated voltage activity. A) Raw voltage correlation matrix from 41 mEC neurons in Fig. 5. B) Agglomerative dendrogram of average voltage correlation linkages from pairwise correlations in A. C) Sorted voltage correlation matrix based on hierarchical clustering optimal leaf order. D) Sorted time series voltage activity based on sorted voltage correlation matrix. E) Spatial organization of voltage correlation clusters across the imaging field of view. F) Pairwise voltage correlations are weakly correlated with distance.

To quantify this potential effect, we used pairwise correlation (r) as a distance metric (d = 1 − r) and performed hierarchical clustering on the voltage correlations. Using an inconsistency coefficient criterion to identify natural groupings, we identified 20 distinct clusters among 41 neurons (Fig. 6B–D). The largest cluster contained 7 neurons, while smaller clusters consisted of only a few or single neurons. Neurons within each cluster exhibited highly similar subthreshold membrane dynamics and firing patterns, indicating shared synaptic input or network drive.

We next mapped cluster identity onto the spatial positions of neurons within the imaging field (Fig. 6E). Clusters of correlated voltage activity were spatially localized, suggesting that subthreshold dynamics are organized within local microdomains. However, this relationship could not be explained by pairwise distance alone, as voltage correlations exhibited only a weak linear dependence on inter-neuronal distance (Fig. 6F). Therefore, spatial organization likely arises from structured connectivity rather than simple proximity.

Voltage signals reflect both subthreshold and spiking activity. To determine whether spatial organization was driven by spike timing, we repeated the clustering analysis using spike trains alone. Spike trains were binarized and smoothed with a Gaussian filter prior to computing pairwise correlations. Compared to voltage signals, spike correlations were lower in magnitude and less variable (Fig. S6A, B), consistent with the fact that all neurons were recruited during theta stimulation. Importantly, clustering based on spike correlations did not reveal clear spatial organization (Fig. S6C, D), and spike correlations showed no relationship with distance (Fig. S6E). Thus, in contrast to subthreshold dynamics, spike timing is spatially distributed across the network.

Together, these results reveal a dissociation between subthreshold and spiking activity. While subthreshold membrane potentials are organized into spatially localized clusters, spike output remains distributed across the network. This suggests that local synaptic inputs and shared network drive are spatially structured, whereas spike generation introduces variability that disperses activity across excitatory populations. Such an organization may enable locally coordinated integration of synaptic inputs while preserving flexible and distributed spiking activity, consistent with models of grid cell coding that incorporate modular organization and distributed spatial phase representations (Barry et al., 2007; Burak and Fiete, 2009; Couey et al., 2013; Hafting et al., 2005; Stensola et al., 2012).

### Computational modeling of CaMKIIα+ network stimulation in layer II/III mEC

In our voltage imaging experiments, we could not simultaneously examine voltage activity in both excitatory neurons and PV+ interneurons. Further, we did not conduct paired patch clamp recordings of excitatory and inhibitory neurons to examine the concurrent timing of synaptic inputs during theta-nested gamma oscillations. To better understand the simultaneous contributions of many excitatory and inhibitory neurons to gamma rhythms, we added 400 heterogeneous biophysically calibrated stellate cells (SCs) to our previously calibrated heterogeneous computational network of 100 PV+ fast-spiking interneurons (Via et al., 2022), as described in the Materials and methods. To our knowledge, this is the first mEC model to capture the full heterogeneity of both excitatory and inhibitory subpopulations. Since only the excitatory cells receive simulated optogenetic drive, as in our experimental studies, computational simulations of gamma oscillations in layer II/III of mEC exhibit the classic PING mechanism (Börgers and Kopell, 2005; Börgers and Walker, 2013; Tiesinga and Sejnowski, 2009), in which PV+ fast-spiking interneurons (PV-INs) fire only when prompted by the excitatory cells.

Figs. 7A1–2 show the simultaneous membrane-potential traces of PV-INs and SCs, respectively, during simulated 8 Hz optogenetic stimulation. The modeling predicted, and experiments validated (see Fig. 1Aiii), the clustering of PV-IN spikes driven by the gamma-synchronous excitatory synaptic inputs into bursts on some gamma cycles, particularly the first cycle. This phenomenon was foreshadowed in Figure 1G of (Pastoll et al., 2013). The model also captured cycle skipping by the stellate cells (compare to Fig. 1Ai); the population rhythm is comprised of different subsets of stellate cells firing on each gamma cycle. Additional examples of both phenomena can be observed in the population raster of the network (Fig. 7A3). The waveforms of the IPSCs received by a voltage-clamped stellate cell model (Fig. 7B1 top) and the EPSCs received by a voltage-clamped PV-IN (Fig. 7B2 top) are qualitatively similar to their experimental counterparts (Fig. 2A, 3A). The fast gamma rhythmicity is clear (Fig. 7B1, B2), and the power is within the experimental range observed (Fig. 2B). The cross correlograms from the experimental data (Fig. 3D) showed that IPSCs and the LFP were highly correlated. Similarly, the average cross correlation correlogram (Fig. 7B3) between the IPSCs (Fig. 7B2) and the EPSCs (Fig. 7B1) shows that the IPSCs follow the EPSCs with a lag of about 2.1 ms, which is the location of the first peak in the cross correlogram, and well within the range of experimental values (Fig. 3G).

**Figure 7:**
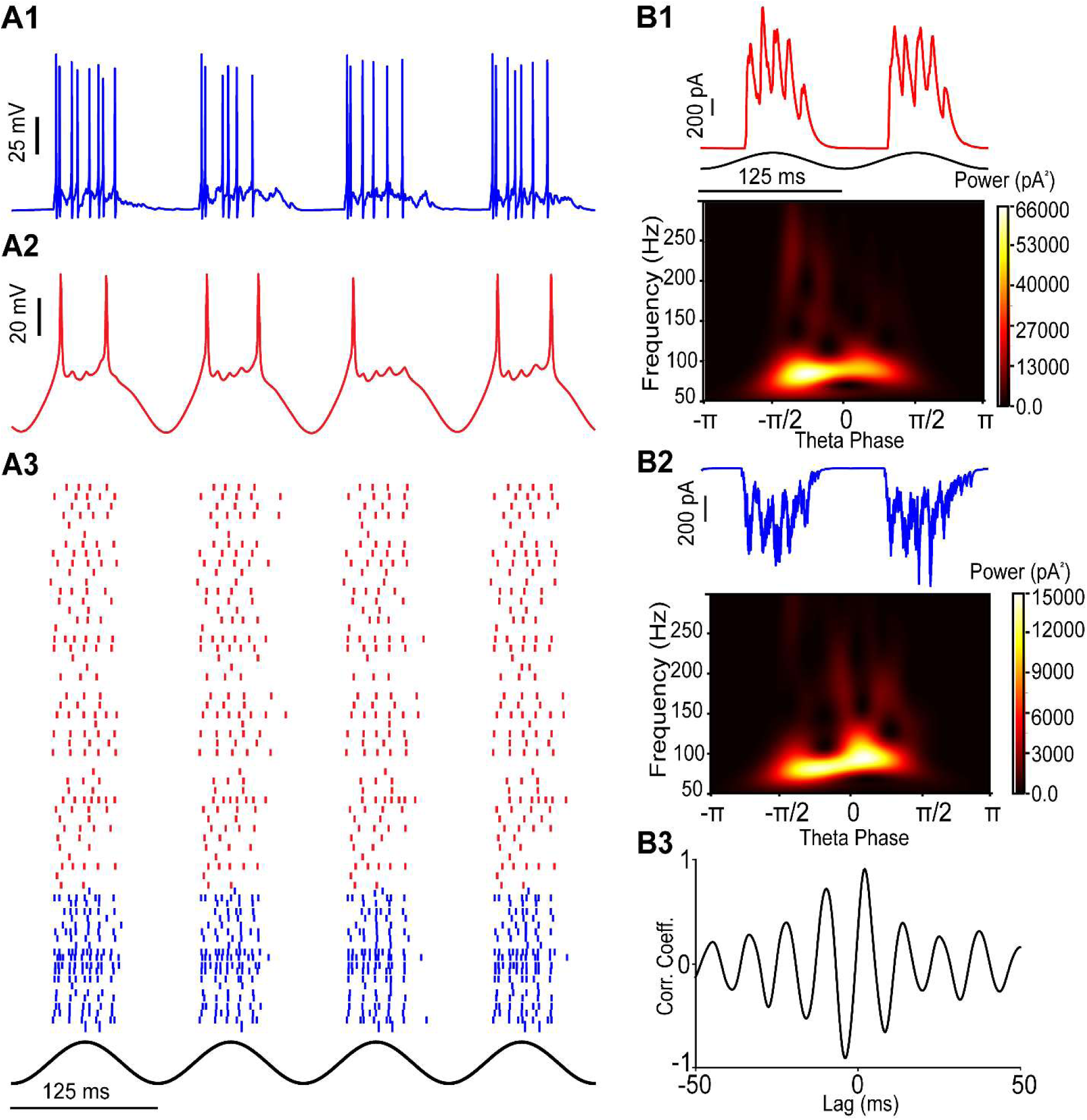
Simulation of PING mechanism during optogenetic theta drive of stellate cells. A1) Voltage traces from PV-INs, A2) Voltage traces from SCs. A3) Downsampled raster plot showing SCs (red) and PV-INs (blue). B1) Top: Inhibitory synaptic currents (red) from a representative SC voltage-clamped at −70 mV. Bottom: Average scalogram of the inhibitory synaptic currents during a theta cycle. B2) Top: Excitatory synaptic currents (blue) from a representative PV-IN clamped at 0 mV. Bottom: Average scalogram of excitatory synaptic currents during a theta cycle. B3) Average cross-correlogram between gamma filtered IPSCs and EPSCs shows that the IPSCs received by the SCs follow the EPSCs recorded in the PV-INs by 3 ms.

## Discussion

In this study, we combined optogenetics, whole-cell patch clamp, large field of view voltage imaging, and LFP recordings in acute CaMKIIα-ChR2 mouse brain slices to investigate the mechanisms underlying theta-nested gamma oscillations in mEC. Our findings strongly support a pyramidal-interneuron network gamma (PING) circuit mechanism in the superficial layers of the medial entorhinal cortex, with gamma oscillations emerging from fast, reciprocal excitation and inhibition between CaMKIIα+ principal cells and fast-spiking interneurons. Optical voltage recordings show evidence of spatial clustering in subthreshold membrane potential, but in this preparation, there is no spatial organization of spike trains.

### PING circuitry generates strong theta-nested gamma oscillations

Our synaptic and firing analyses revealed that fast-spiking interneurons receive strong gamma-frequency excitatory drive, sometimes firing in rhythmic bursts, and provide potent feedback inhibition onto principal cells. This supports our previous prediction that excitatory cell input can group PV+ firing into bursts as a means of slowing their contribution to the oscillations recorded in the LFP (Williams et al., 2026). Notably, gamma-frequency EPSCs were nearly absent in stellate and pyramidal cells, while IPSCs were both strong and gamma-rhythmic. This temporal structure, where EPSCs in interneurons precede IPSCs in principal neurons by ∼3 ms, along with phase-locking of IPSCs to the LFP gamma, provides strong evidence for a feedforward excitation followed by feedback inhibition motif, supporting a PING mechanism in mEC (Butler et al., 2018; Pastoll et al., 2013). The absence of recurrent excitation is consistent with connectivity studies (Couey et al., 2013; Dhillon and Jones, 2000; Fuchs et al., 2016; Pastoll et al., 2013), besides one (Winterer et al., 2017), and supports grid cell models that connect principal neurons exclusively through feedback inhibition. Moreover, the IPSCs in excitatory cells appear to be very similar to the recorded LFPs, potentially dispensing with the need for complex models of the LFP (Bédard and Destexhe, 2012; Leung, 2011; Lindén et al., 2014).

Our voltage imaging results further confirm that excitatory neurons exhibit gamma-synchronous spiking across the network, despite mostly slower interspike rates at the level of individual cells. This supports a model where different populations of excitatory neurons fire on alternating gamma cycles, maintaining coherent network-level oscillations while individual neurons skip cycles (Buzsáki and Wang, 2012; Pastoll et al., 2013).

### ING circuitry cooperatively synchronizes gamma oscillations

While optogenetic activation of CaMKIIα+ neurons drive a PING circuit mechanism, fast-spiking interneurons receive gamma frequency IPSCs, indicating that recurrent inhibition plays a cooperative role in shaping theta-nested gamma oscillations in mEC. Although the power of IPSCs in fast-spiking interneurons are significantly weaker than those received by principal neurons, the inhibitory-inhibitory (I-I) feedback loop may further facilitate synchronized activity by turning off the strong inhibition onto principal neurons. In addition to chemical synapses, PV+ putative fast-spiking interneurons are interconnected via electrical synapses, or gap junctions (Fernandez et al., 2022; Huang et al., 2024). These electrical connections allow for rapid and bidirectional communication between interneurons, promoting synchronous firing and enhancing the precision of network oscillations. Although I-I feedback loops are not necessary for generating models of gamma oscillations (Börgers and Kopell, 2003; Brunel and Wang, 2003), the addition of I-I connectivity enhances grid stability and increases gamma frequency (Shipston-Sharman et al., 2016; Solanka et al., 2015). Our results suggest that E-I and I-I hybrid gamma networks may work together to generate gamma frequency oscillations.

### Cell-type specific differences in gamma phase locking

Stellate cells, pyramidal cells, and fast-spiking interneurons exhibited strong theta phase locking, confirming entrainment to optogenetic stimulation. At the single-cell level, gamma phase locking was comparable across cell types, but spike timing was systematically offset. Pyramidal neurons fired earlier in the gamma cycle, predominantly along the descending phase, whereas stellate cells and fast-spiking interneurons fired near the trough. The phase difference between pyramidal cells and fast-spiking interneurons corresponded to ∼3 ms, closely matching the delay between excitation and inhibition measured in interneurons and supporting a PING mechanism. Stellate cells appear to follow the firing of pyramidal cells and play a supporting role in the recruitment of local interneurons.

A previous model of grid cell activity predicted gamma phase locking in both excitatory and inhibitory neurons (Pastoll et al., 2013), but experimental recordings showed weak gamma locking at the population level for stellate cells and fast spiking interneurons. In contrast, our results demonstrate that both individual neurons and population analysis of all cell types exhibit gamma phase locking. Notably, pyramidal cells were more consistently engaged with gamma oscillations than stellate cells, a distinction that has not been fully explored in earlier work.

These findings suggest that pyramidal cells may play a more important role in the gamma generating microcircuit than previously recognized. The current study did not model two groups of excitatory cells but focused instead on stellate cells. Future models should elucidate any differential contributions of the two subtypes.

### Comparison to previous models of theta-nested gamma in the mEC

Previously (Williams et al., 2026), we modeled optogenetically-driven theta nested gamma in an mEC mouse slice under different conditions. The ChR2 was expressed using the Thy1 promotor rather than the CaMKIIα promotor used here. Therefore both excitatory and inhibitory neurons received strong theta drive (Pastoll et al., 2013), whereas in this study the optogenetic drive was mostly confined to the excitatory population (Butler et al., 2018). The major difference from our previous study is that in models with a significant drive to the I cells, they do not require input from the E cells to fire. Thus in order for the E cells to control the frequency of the I cells, the drive to the E cells must be sufficiently strong so that they recover from inhibition before the I cells do, and the connection strength of the E to I synapses must be sufficiently strong to slow the ING frequency into a PING range, by increasing total inhibition in the network (sometimes by evoking bursts in the I cells). We call this an “E cell recovers first” PING mechanism (Whittington et al., 2000). In contrast, models with low to no I cell drive exhibit a classic PING mechanism (Börgers and Kopell, 2005; Börgers and Walker, 2013; Tiesinga and Sejnowski, 2009) in which the I cell firing is evoked by a population burst in the E cells, and the I cells silence firing between E cell population bursts. We call this mechanism a “driven I cell” PING. This study, like previous studies (Pastoll et al., 2013; Solanka et al., 2015; Sutton et al., 2024) utilizing less biophysically realistic neurons, employs the latter mechanism.

Another important difference from our previous model of PING in the mEC is that it used generic Hodgkin-Huxley model neurons for the E cell rather than biophysically calibrated stellate cell models. The theta drive was sufficient to synchronize the weakly heterogeneous stellate cells in that model (Williams et al., 2026), therefore external noise had to be added to each E cell to prevent hypersynchrony. Because the current study captured the full range of heterogeneity, the addition of noise was no longer necessary. Our computational model is the most biophysically realistic model to date of theta-nested gamma in the mEC because both the inhibitory and excitatory populations have been carefully calibrated to capture the characteristic electrophysiological properties of the respective populations plus the full range of heterogeneity in those properties. The model also captures the spiking patterns of both inhibitory and excitatory cells exhibited during theta drive. Future studies will incorporate this network into a model of grid cell activity by incorporating velocity inputs tuned to a specific movement direction as in (Couey et al., 2013; Pastoll et al., 2013) to improve our understanding of how activity bumps and make predictions regarding single versus multiple bump attractors (Shipston-Sharman et al., 2016).

### Voltage imaging captures population-level synchrony and correlated subthreshold activity in spatial clusters of principal neurons

Large field-of-view voltage imaging allowed us to simultaneously record from over 40 neurons, providing an unprecedented mesoscopic view of intracellular voltage activity. We observed network-wide gamma frequency synchronization over distances greater than 500 microns across layer II mEC. This suggests that local connectivity is sufficient to coordinate gamma-synchronous firing across hundreds of microns, supporting the idea that gamma oscillations reflect global coordination within a modular microcircuit (Buzsáki and Wang, 2012). Figure 5E shows the first population spike raster plot for excitatory cells during theta nested gamma oscillations. Previous work on optogenetically driven theta nested gamma in the mEC focused on the LFP (Butler et al., 2018) and spike histograms on repeated trials in single neurons (Pastoll et al., 2013), but the population histograms in this study are novel, and better reveal the cycle skipping nature of the participation of the excitatory cells.

Spike correlations revealed no linear distance dependence or spatial clustering, while voltage correlations demonstrated a weak pairwise distance relationship and local clusters of activity. Therefore, the subthreshold activity or spike shape of nearby excitatory neurons is likely driving the correlated spatial patterns. The laminar structure of the mEC may also influence the spatial organization of neural activity. For example, neurons in layer III appear to be much less correlated to neurons in layer II. Our fields of view were primarily centered on layer II and limited to 300 microns across cortical layers, it’s likely that we could not capture the simultaneous dynamics of both layer II and III. Additionally, the sparse connectivity of principal cells in layer II (especially stellate cells) may underlie the distributed spike correlations.

Interestingly, we found that increasing the frequency of theta stimulation decreased the number of gamma cycles per theta period without changing gamma frequency itself. This aligns with predictions from PING models where gamma frequency is determined by local E/I kinetics, while theta pacing controls the envelope (Traub et al., 1996; Wang, 2010).

### Limitations and technical considerations

Although our voltage imaging experiments provided an unprecedented view of network dynamics in the mEC, several technical limitations constrained the scope of our findings. First, immunohistochemical analysis revealed that parvalbumin-positive (PV+) interneurons did not express the genetically encoded voltage sensor Voltron2, limiting our recordings primarily to putative excitatory neurons. This lack of PV+ labeling is likely due to the poor tropism of the AAV2retro serotype for PV+ interneurons in the mEC. While a small fraction of other interneuron subtypes may be present in the dataset, their contribution is likely minimal.

Second, imaging with a high-powered 561 nm laser, used to excite Voltron2-JF585, induced some level of ChR2-mediated current. This led to mild depolarization and spontaneous firing in CaMKIIα+ neurons prior to the onset of widefield optogenetic stimulation (Fig. 5B). However, this effect was transient: neuronal excitability normalized shortly after the start of rhythmic stimulation, as evidenced by stable firing during subsequent theta cycles and the return to baseline activity following stimulation. Importantly, the LFP gamma oscillations and excitatory neuron firing rates recorded during voltage imaging were consistent with those observed in parallel patch-clamp experiments.

Third, the 800 Hz imaging frame rate, combined with the temporal filtering properties of the Voltron2 sensor, limited our ability to resolve gamma phase-locking at the single-cell level. As a result, detailed analyses of spatial gamma phase relationships were not feasible.

### Future directions

Future work should employ *in vivo* voltage imaging or multiphoton imaging of genetically defined subtypes (e.g., stellate (reelin+), pyramidal (calbindin+), fast-spiking interneurons (PV+)) to clarify their roles in spatial coding during behavior and theta-nested gamma oscillations.

Additionally, closed-loop optogenetic perturbations of fast-spiking interneurons could test causality in gamma generation in mEC. Computational modeling that incorporates our measured E-I delays, phase-locking dynamics, and correlated spatial clusters may further refine existing models of theta-gamma nesting and grid formation.

## Materials and methods

All experimental procedures were conducted using Boston University IACUC-approved protocols. Adult (2-6 months old) male and female mice were used in roughly equal numbers. A transgenic mouse line expressed Cre recombinase under the CaMKIIα promoter (CaMKIIα-Cre, JAX strain #005359). CaMKIIα is expressed in mainly excitatory neurons in mEC. The transgenic CaMKIIα-Cre mouse line was cross bred with transgenic LoxP-ChR2-EYFP mice (JAX, strain #024109) to constitutively express ChR2 and EYFP in CaMKIIα+ neurons (CaMKIIα-ChR2-EYFP) after Cre-Lox recombination.

### Viral injection procedure

Adult (2-6 months old) transgenic CamKIIα-ChR2 mice were intracranially injected with AAV2retro-Syn1-Voltron2-ST-WPRE-BGHpA (Titer: 1.2 x 10^12^ GC/mL, custom plasmid, UNC NeuroTools Vector Core) at least three weeks prior to voltage imaging experiments. Viral injections were unilateral and alternated between right and left hemisphere for each mouse. Mice were weighed prior to surgery to set the proper flow rate for the nose cone and dosage for Buprenorphine administration. Mice were induced with 3.0% isoflurane and moved to a nose cone with maintenance anesthesia gradually reduced from 2.3% to 1.7% throughout the surgery. Body temperature was maintained using a heating pad set to 37.2°C. A craniotomy was drilled above the desired stereotaxic location and a nanofil syringe delivered virus to two different locations across the dorsoventral axis of the mEC (−4.90 A/P, ±3.20 M/L, −3.50 D/V, 500 nL; −4.60 A/P, ±3.20 M/L, −4.00 D/V, 500 nL; from bregma).The virus is injected at 75 nL/min and removed between injections. After injections, the skin was sutured with Vetbond and Buprenorphine was administered prior to removal from anesthesia. Mice were given 6 doses of Buprenorphine (.05 mg/kg), once every 12 hours, starting prior to removal from anesthesia. Diet supplements and hydrogels were given to aid recovery.

### Acute slice preparation

All whole-cell electrophysiology measurements were obtained from 400-micron thick horizontal mouse brain slices, while 300-micron thick slices were used to maximize the viral expression at the imaging plane for voltage imaging experiments. Bilateral slices from the dorsomedial mEC (∼3.2-4.3 mm from the dorsal surface of the brain) were used. Mouse brains were extracted after isofluorane overdose and decapitation. During slicing, the brain was submerged in sucrose-substituted artificial cerebrospinal fluid (ACSF) solution (in mM: sucrose 225, KCl 2.5, NaH_2_PO_4_ 1.25, MgCl_2_ 3, NaHCO_3_ 25, glucose 20, and CaCl_2_ 0.5) at 4°C that was continuously perfused with 95% oxygen / 5% carbon dioxide (carboxy) gas and sliced with a vibratome (VT1200, Leica Microsystems). Then, brain slices are moved to a separate chamber containing standard ASCF (in mM: NaCl 125, NaHCO_3_ 25, D-glucose 25, KCl 2.5, CaCl_2_ 2, NaH_2_PO_4_ 1.25, and MgCl_2_ 1) perfused with carboxy gas and incubated at 37°C for 30 minutes. After incubation, the brain slices are allowed to recover for 15 minutes at room temperature (∼20°C) where they remained until being used for whole-cell electrophysiology. For voltage imaging experiments, brain slices were moved to secondary carboxy-perfused incubation chamber containing 5 mL ACSF and 25 nmol of JF585 Halotag ligand dye for 1 hour. The JF585 dye was dissolved in 10 μL dimethyl sulfoxide (DMSO) before mixing with the ACSF. Following the 1-hour incubation, brain slices were moved back to the primary 200 mL ACSF chamber to washout unbound JF585 dye for 30 minutes prior to imaging.

### Acute slice electrophysiology

For all electrophysiology experiments, brain slices were continuously perfused with ACSF at a temperature of 35–37°C. The ACSF was bubbled with 95%/5% carboxy gas throughout the experiments. Electrophysiology data was amplified using an Axon Instruments MultiClamp 700B and sampled using Axon Instruments DigiData 1440A at 30 kHz for whole-cell experiments or 33 kHz for voltage imaging experiments. Custom protocols were designed using pClamp 7.0 software to control data collection and optogenetic stimulation. Electrodes were pulled using a horizontal puller (Sutter Instrument) using filament, thin-wall borosilicate glass (Sutter Instrument). Extracellular pipettes were filled with ACSF and pipettes with access resistances of 0.5–2 MΩ were inserted into layer II/III mEC for simultaneous LFP recordings. Extracellular recordings were hardware filtered from 0.1–1000 Hz.

For whole-cell recordings, borosilicate glass pipettes were filled with intracellular fluid solution (in mM: K-gluconate 136, KCl 4, HEPES 10, diTrisPhCr 7, Na_2_ATP 4, MgCl_2_ 2, TrisGTP 0.3, and EGTA 0.2, and buffered to pH 7.3 with KOH) and pipettes with resistances of 4-8 MΩ were used. Pipette offset was compensated prior to achieving on-cell patch recordings. Once the patch electrode was sealed to the cell membrane (>1 GΩ seal), pipette capacitance was compensated. Series resistances < 40 MΩ were used in this study with changes < 20% throughout recordings. For voltage clamp recordings, series resistance was compensated 50–70%. Using voltage clamp, excitatory and inhibitory post-synaptic currents were recorded at −70 and 0 mV, respectively. For our recording conditions, the reversal potential for chloride is −75 mV. For current clamp recordings, full bridge balance compensation was used. Liquid junction potentials were not corrected. Pipettes and cells were visualized with diffuse interference contrast.

### Optogenetic stimulation

During whole-cell recordings, a 470 nm LED (Thorlabs, M470L4) delivered widefield sinusoidal optogenetic stimulation through a 40x objective lens. For each cell, the same light stimulation power was used throughout recordings (i.e., voltage clamp at −70 mV and 0 mV; current clamp at 0 pA). For compatibility with the voltage imaging microscope, a 490 nm LED (Thorlabs, M490L4) delivered widefield sinusoidal optogenetic stimulation through a 16x objective lens. For each field-of-view, the same light stimulation power was used throughout recordings (i.e., 4, 8, 12, or 16 Hz sinusoidal stimulation). However, the light power was adjusted for each cell (field-of-view) to drive sufficient activity for gamma frequency synchronization in the surrounding local network similar our prior study (Williams et al., 2026). Previously, we found no relationship between optogenetic stimulation intensity and the frequency or power of gamma oscillations in layer II/III mEC across different sessions in Thy1-ChR2 mice. The optimal light power varied to correct possible variations in slice excitability and functional connectivity.

### Electrophysiological cell type classification

For whole-cell recordings, the major electrophysiological cell types in the mEC were statistically separated by their electrophysiological properties. Current steps were injected from −200 pA to 525 pA at 25 pA intervals to characterize the subthreshold and firing properties of each cell.

Putative stellate and pyramidal cells in layer II/III mEC were classified based on membrane sag potential and membrane time constant as used previously (Fernandez et al., 2022; Williams et al., 2026). Fast-spiking interneurons were easily identified by their discontinuous membrane current – spike rate relationship, high threshold firing, burst firing, fast membrane time constant, and short spike half width.

### Targeted illumination confocal voltage imaging and optogenetic stimulation

A line-scan, targeted-illumination confocal microscope (Xiao et al., 2024) was used to image the chemigenetic voltage indicator, Voltron2-JF585, with high signal to noise ratio over a large field of view. At 800 Hz frame rate, we recorded from a large field of view up to 1.16 × 0.325 mm using a 16x objective lens (Nikon ×16/0.8 NA LWD). A 561 nm laser (Oxxius, LCX-561L-200-CBS-PPA; 200 mW output power) was used for Voltron2 imaging and a digital micromirror device (DMD) enabled targeted illumination. A combination of excitation filter, emission filter and dichromatic mirrors (Chroma Technology Corp., 89901v2; 405/488/561/640-nm Laser Quad Band Set) separated fluorescence from the excitation light. A 490 nm LED (Thorlabs, M490L4) and 488 nm bandpass filter (Edmund Optics, 65-147) were added to the excitation path for widefield optogenetic stimulation through the 16x objective lens. A 575 nm long pass emission filter (Chroma, ET575lp) was inserted prior to the sCMOS camera (Teledyne Photometric Kinetix) to block both optogenetic excitation light and EYFP emission.

### Voltage imaging data acquisition

Teledyne Photometrics PVCAM software was used for image acquisition. Reference images were acquired in TIFF format and a custom graphical user interface (MATLAB) enabled users to draw rectangular regions of interest for targeted illumination. All video recordings were acquired in RAW format prior to data analysis. Custom protocols were written using pClamp 7.0 software to control simultaneous optogenetic stimulation, voltage imaging, and LFP recordings. Several different sinusoidal frequencies were used for optogenetic stimulation: 4, 8, 12, and 16 Hz. Each video recording started and ended with 500 ms of baseline activity without stimulation. The number of stimulation periods varied for each simulation frequency: 16 cycles for 4 Hz, 23 cycles for 8 Hz, 17 cycles for 12 Hz, 16 cycles for 16 Hz. The same laser imaging intensity was used for all recordings in a single field of view. Laser imaging intensities varied from 50–70 mW/mm^2^ across recording sessions.

### Immunohistochemistry

Immunohistochemistry was performed to identify parvalbumin-expressing interneurons in acute voltage imaging experiments. Acute mouse brain slices were fixed in 4% paraformaldehyde (PFA) immediately after voltage imaging experiments and stored in the 4°C fridge overnight.

Approximately 16 hours later, slices were moved into wells containing phosphate-buffered saline (PBS) solution and placed on a shaker at room temperature for 15 minutes. The wells were swapped with fresh PBS solution and washed two more times. Slices were temporarily stored in PBS-NaN_3_ (1% sodium azide) if not immediately proceeding with immunohistology. First, slices were blocked with 5% normal goat serum (Thermo Fischer Scientific) and 95% PBST [PBS with 0.2% v/v TritonX (Thermo Fischer Scientific)] for 1 hour at room temperature. Slices were then moved to wells containing the primary antibodies, rabbit anti-parvalbumin (Swant PV 27), 1:1000 in PBST and 2% normal goat serum, and chicken anti-GFP (Thermo Fischer Scientific), 1:1000 in PBST and 2% normal goat serum. The primary antibodies were incubated on a shaker at room temperature overnight. The following day, slices were washed three times with PBST on a shaker at room temperature for 15 minutes each. After washing, slices were moved to wells containing the secondary antibodies, Alexa Fluor 647 goat anti-rabbit, 1:500 in PBST and 2% normal goat serum, and Alexa Fluor 488 goat anti-chicken, 1:200 in PBST and 2% normal goat serum. The secondary antibodies were incubated on a shaker for 4 hours at room temperature.

After secondary antibody incubation, slices were washed three times with PBST on a shaker for 15 minutes each at room temperature. Slices were then mounted on a slide using Vectashield with DAPI (Vector Laboratories) and a coverslip. Slides were left in a 4°C fridge overnight prior to imaging.

### Confocal imaging and post-hoc image registration

Slides were imaged on an Olympus FV3000 confocal microscope using a 20x air objective. A stack of images was acquired at 5 μm intervals for each slice. Neurons that expressed both Parvalbumin and Voltron2-JF585 were manually registered to the corresponding TICO scope reference image to identify active PV+ interneurons during optogenetic stimulation. Confocal image stacks and TICO reference images were background subtracted, and contrast enhanced using ImageJ.

### Voltage trace extraction and spike detection

Video data was extracted from RAW files and regions of interest (ROIs) were manually drawn for each neuron. The raw fluorescence (𝐹_raw_) was calculated as the sum of the pixel intensities within each ROI for each frame. Motion was minimal due to the fixed nature of the slices and short recordings. Additionally, photobleaching was minimized as less laser power is needed to image neurons at a short depth (50-100 µm) below the surface of acute brain slices. Nonetheless, slow changes in fluorescence were filtered out prior to spike detection, 𝐹_fast_.The raw fluorescence, 𝐹_raw_, was forward and reverse filtered with a 4^th^ order bandstop Butterworth filter (bandstop cutoff: 0.1-3 Hz) to obtain 𝐹_fast_. The change in fluorescence was calculated by subtracting the mean fluorescence of 𝐹_fast_, 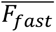 and then dividing by the mean fluorescence, 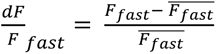. Since Voltron2 is a negative-going voltage indicator, meaning that neural depolarizations are reflected as negative changes in fluorescence, 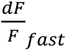 was inverted, 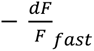. The noise in each trace was estimated using 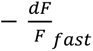 for the first 100 ms after optogenetic stimulation was turned off, 𝐹_noise_, as neurons were least likely to fire action potentials during this period. First, 𝐹_noise_ was detrended, then 𝐹_noise_ was limited to negative changes (𝐹_noise_ < 0) to reduce spike contamination. Noise (𝜎_noise_) was estimated as the standard deviation of the negatively rectified, detrended fluorescence trace.

For spike detection, 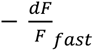 was forward and reverse filtered with a 4^th^ order high pass Butterworth filter (high pass cutoff: 50 Hz) to remove subthreshold activity 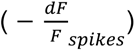. The *findpeaks* function in MATLAB was used to detect spikes in 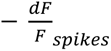. The following parameters were empirically found to detect spikes with high fidelity: 𝑀𝑖𝑛𝑃𝑒𝑎𝑘𝐷𝑖𝑠𝑡𝑎𝑛𝑐𝑒: 2 𝑠𝑎𝑚𝑝𝑙𝑒𝑠 (2.5 µ𝑠), 𝑀𝑎𝑥𝑃𝑒𝑎𝑘𝑊𝑖𝑑𝑡ℎ: 6 𝑠𝑎𝑚𝑝𝑙𝑒𝑠 (7.5 µ𝑠), 𝑀𝑖𝑛𝑃𝑒𝑎𝑘𝑃𝑟𝑜𝑚𝑖𝑛𝑒𝑛𝑐𝑒: 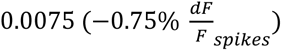, 𝑀𝑖𝑛𝑃𝑒𝑎𝑘𝐻𝑒𝑖𝑔ℎ𝑡: 2𝜎_noise_, and 𝑊𝑖𝑑𝑡ℎ 𝑅𝑒𝑓𝑒𝑟𝑒𝑛𝑐𝑒: ℎ𝑎𝑙𝑓ℎ𝑒𝑖𝑔ℎ𝑡.

After spike detection, spikes were removed from 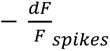 and the standard deviation of the spike removed (SR) data was calculated as an updated estimation of noise (𝜎_SR noise_). Spikes detected with peak heights less than 3.5 standard deviations of the updated noise estimate were removed from analysis (𝑝𝑒𝑎𝑘𝑠 < 3.5𝜎_SR noise_). The signal to noise ratio (SNR) of each neuron was calculated as average spike peak height divided by the updated standard deviation of noise 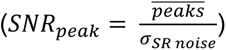. Neurons with 𝑆𝑁𝑅_peak_ less than 3.75 were removed from analysis. Furthermore, neurons that were constantly firing action potentials were removed from analysis. All traces were manually inspected to ensure that spike fidelity was greater than 90%.

### Whole-cell current analysis

For current recordings, raw data was forward and reversed filtered from 50–200 Hz with 4th order butterworth filters. To analyze the frequency and amplitude of gamma oscillations, the continuous wavelet transform was computed in MATLAB. The analytic morlet wavelet (ω_0_ = 6 rad/sec) was used to calculate the scalogram for each theta cycle with 32 scales per octave. The network activity during the first stimulation period varied greatly compared to subsequent cycles and was removed from analysis. The magnitude of the scalogram was averaged over the next 40 theta cycles to determine the peak gamma frequency, phase and power of the membrane currents for each cell. Any theta cycles with large current artifacts or spikes (> 3000 pA) were removed from analysis. Low average peak gamma power recordings (< 20 pA^2^), scalograms peaks with large bandwidths (> 100 Hz), and scalograms with peak powers less than five multiples above the average (SNR < 5) were removed for accurate gamma frequency comparisons.

For whole-cell current and LFP cross correlation analysis, raw data was forward and reverse filtered from 50–200 Hz with 4th order butterworth filters. For each recording, gamma-filtered current and LFP data was cross-correlated and averaged across stimulation cycles 2–6. The peak correlation coefficient was positive for inhibitory currents and negative for excitatory currents. The absolute value of the peak correlation coefficient was reported in Fig. 3E, F. The lag of the peak correlation coefficient indicates a negative value if the synaptic currents precede the LFP.

### Local field potential analysis

Raw local field potential recordings were filtered in both theta and gamma frequency ranges. The bandpass cutoff for theta range filtering was adjusted based on the stimulation frequency (4 Hz stim: 2–6 Hz; 8 Hz stim: 4–12 Hz; 12 Hz stim: 8–16 Hz; 16 Hz stim: 12–20 Hz). The bandpass for gamma frequency filtering was dependent on the analysis. For frequency spectrum and cross correlation analysis, the 50–200 Hz range was used. All filters were 4th order Butterworth bandpass filters and applied in the forward and reverse direction. The analytic morlet wavelet (ω0 = 6 rad/sec) was used to calculate the scalogram for each theta cycle with 32 scales per octave. For voltage imaging and LFP recordings, LFP scalogram power was averaged all theta cycles to determine the peak gamma frequency, phase and power of the LFP. Any theta cycles with large voltage artifacts (> 1000 µV) were removed from analysis. To determine the phase of the gamma oscillations in the LFP using the Hilbert transform, a more restrictive bandpass filter is necessary to ensure that the signal is nearly a single frequency sinusoid prior to phase estimation. For the voltage imaging and LFP phase locking analysis, 60–120 Hz was used. For the intracellular patch clamp voltage recordings and LFP phase analysis, 60–120 Hz was used.

### Spike – gamma phase analysis

The instantaneous phase of the gamma-filtered LFP components was obtained using the Hilbert transform. For whole-cell voltage and LFP recordings, action potential peaks from the first 40 theta stimulation cycles were registered to the phase of the gamma-filtered (60–120 Hz) LFP data. If a spike burst was detected (interspike intervals < 6.67 ms), only the first spike phase was used for phase analysis. For simultaneous voltage imaging and LFP experiments, the instantaneous phase and filtered traces were down sampled to the imaging frame rate (800 Hz). The unwrapped phase of the LFP components was linearly interpolated for down sampling, then wrapped again from −π to π, while the filtered traces were down sampled using a cubic interpolation method. Action potential peaks were registered to gamma LFP phases for spike phase analysis. The unbiased phase locking value was calculated using the pairwise phase consistency method (PPC; (Vinck et al., 2010)) to evaluate the consistency of phase-locking in individual neurons across theta cycle spikes. Vector strength (i.e., phase locking value) was used to evaluate the phase locking among both individual neurons and cell type populations. Spike phase polar plots used 36 equally spaced bins (10 degrees per bin) to display population gamma locking.

### Spike – theta phase analysis

Action potential peaks were detected and registered to the phase of the theta stimulation period. Spike phase histograms were computed with 50 equally spaced bins over the stimulation period 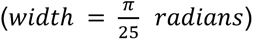 and were normalized by the number of theta cycles (40 cycles for each cell). Interspike frequency histograms were computed with 10 Hz bins and the bin counts were normalized by the number of theta cycles (40 stimulation periods for each cell).

### Spatial correlation analysis

The binary spike mask for each neuron during sinusoidal stimulation was convolved with a normalized gaussian filter kernel [σ = 2 samples (2.5 ms), total size = 6σ] to reduce variability. The correlation coefficient (𝑅_spike_) was calculated between every neuron pair in a field of view. Pairwise distance was calculated as the Euclidean distance between the pixel centroids of each neuron.

### Hierarchical clustering of neural ensembles

We chose an inconsistency coefficient threshold of 1.1 standard deviations above the mean height of links within two levels below it. The dendrogram, correlation matrix and time series data were sorted based on the optimal leaf order to visualize similarities.

### Data analysis and statistics

All data analysis was performed with custom written algorithms in MATLAB. Normality was determined by the Shapiro-Wilk test at a significance of p < 0.05. Levene’s test was used to assess the equality of variances at a significance of p < 0.05. For statistical comparisons between normally distributed with equal variances data, data was reported as mean ± S.E.M. or mean (95% confidence interval) and one-way ANOVA test was evaluated for a significant difference at p < 0.05. If a significant difference was found, the Tukey-Kramer test was applied for multiple comparisons. Spectral power was log-transformed prior to statistical analysis. For non-normal or skewed distributions, data was specifically reported as median (1^st^ quartile, 3^rd^ quartile) and the one-way Kruskal-Wallis ANOVA test determined a significant difference at p < 0.05. If significance was detected, Dunn’s test with Šidák correction was used for multiple comparisons.

### Computational methods

The computational modeling network was comprised of 500 single-compartment neurons of two types: 100 inhibitory PV+ interneurons (I cells) and 400 excitatory stellate cells (E cells). We limited the model to a single population of excitatory cells and thus did not incorporate pyramidal cells at this time. Network simulations were carried out using the NetPyNE (Dura-Bernal et al., 2019) simulator based on Python. Associated programs to run simulations can be found on https://github.com/riyadahal/mEC_PING. Optogenetic theta drive was simulated as a half-wave-rectified 8 Hz sinusoidal conductance waveform multiplied by a driving force calculated with a reversal potential of 0 mV to produce a current drive. The theta drive had a peak conductance of 15 nS and was applied only to the stellate cells, since CAMKIIα is mostly expressed in the excitatory cells. During the theta stimulation of the E cells, we recorded inhibitory and excitatory currents from five readout neurons from each population voltage-clamped at 0 mV and –70 mV, respectively. Oscillation frequency and power in the inhibitory currents were analyzed using the analytic Morlet wavelet. The network model and the Morlet wavelet, as implemented in NeuroDSP (Python) package, were used to generate time-frequency scalograms.

#### PV+ interneuron model

In our previous work (Via et al., 2022), we calibrated 100 PV+ interneurons using data based on the passive and intrinsic properties of these neurons (Fernandez et al., 2022). We use the same model for the interneurons in this work.

#### Excitatory stellate cell model

We used a previously published heterogeneous population of single-compartment stellate cell models from (Mittal and Narayanan, 2018), who conducted a large stochastic parameter search by varying 55 parameters to capture intrinsic heterogeneity in the entorhinal SCs. They identified 449 valid models that satisfied electrophysiological criteria for layer II mEC stellate cells. From this population, we selected 400 stellate cell models for network simulations. Since the stellate cell models contained a slow process, activation, and deactivation of the H current as well as Ca^2+^ dynamics, all simulations were run without any stimulation for 5 seconds to eliminate transient dynamics.

#### Connectivity

The probability of I->I, I->E, and E->I connections were 35%, 20%, and 25%, respectively. I->E synaptic weights were drawn from a lognormal distribution as in (Fernandez et al., 2022) but scaling the original mean by a factor of 5. Gap junctions were included for the I cells with a probability of 20% connection for each pair. In agreement with data from previous experimental studies, the stellate cells were not connected to each other (Couey et al., 2013; Dhillon and Jones, 2000; Fuchs et al., 2016; Pastoll et al., 2013; Williams et al., 2026).

#### Synapses

Both electrical and chemical synapses were modeled for the I->I connections based on our previous work (Via et al., 2022). The I->I inhibitory synapses were calibrated with a rise time constant and decay time constant of 0.3 ms and 2 ms, respectively, with a synaptic reversal potential of E_GABA_ = −75 mV for the chemical synapses. Inhibitory synapses onto the excitatory cells were calibrated with a rise time constant and decay time constant of 0.4 ms and 6 ms, with E_GABA_ = −65 mV (Sauer et al., 2012). The synaptic connections from E cells to the I cells are modeled using an exponential synapse with a decay time constant of 1 ms, a reversal potential of E_AMPA_ = 0 mV, and peak conductance of 0.8 nS. We used randomized synaptic delay to model all chemical synapses following a uniform distribution of 0.6–1 ms.

## Acknowledgements

This work was supported by NIH R01NS054281 to C.C.C. and J.A.W., NIH F31NS134309 to B.W., NSF Grant No. 2018936 to C.C.C., and NIH RF1MH126882 to M.N.E.

## Supporting Information

**Figure S1:**
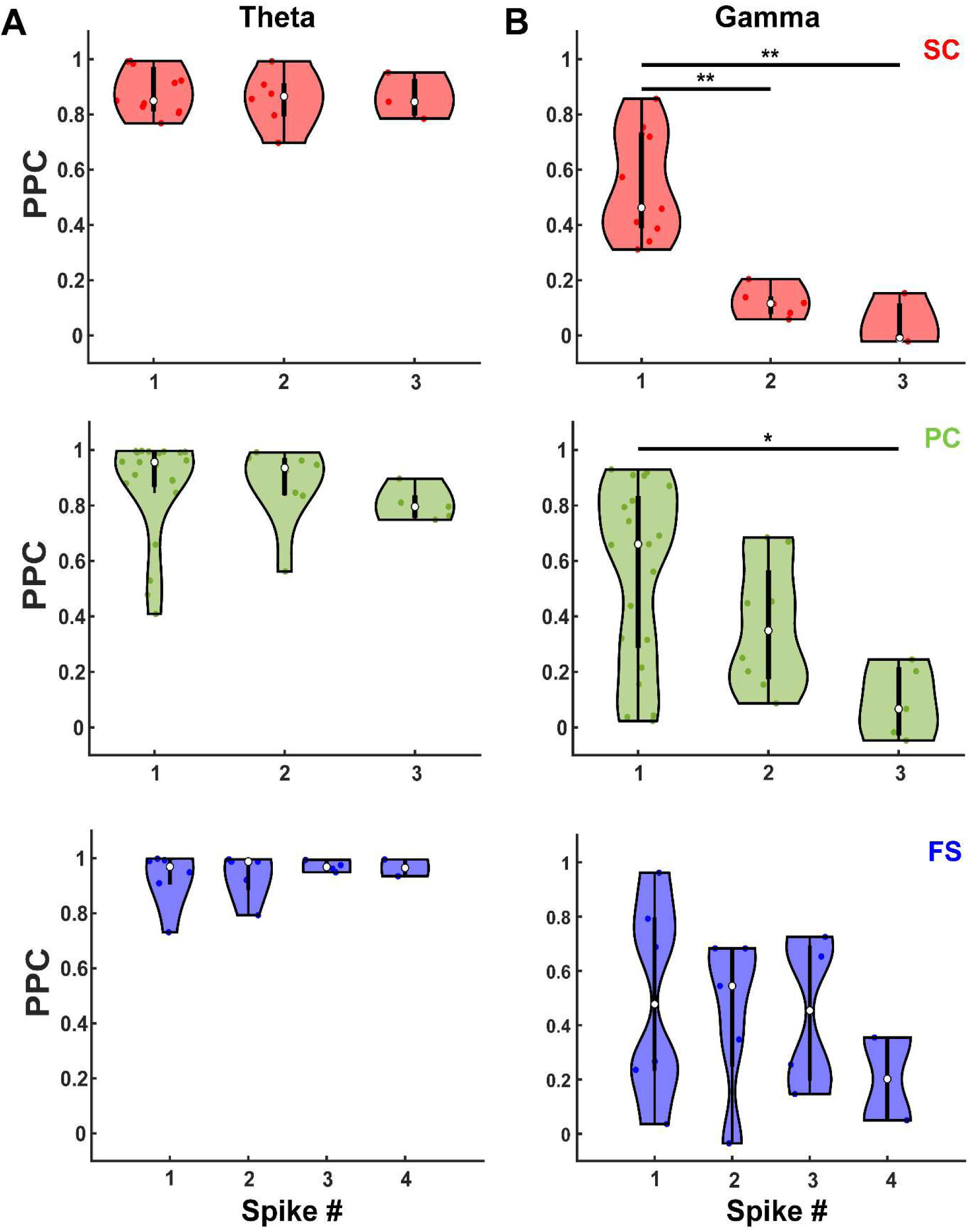
mEC neurons are strongly phase locked to theta and moderately phase locked to gamma oscillations. A) Pairwise theta phase consistency of stellate (red), pyramidal (green), and fast-spiking interneurons (blue) for each consecutive spike per theta stimulation period. All cell types and spike numbers are strongly phase locked to theta drive. B) Pairwise gamma phase consistency of stellate (red), pyramidal (green), and fast-spiking interneurons (blue) for each consecutive spike per theta stimulation period. The first spike per theta stimulation period is moderately phase locked to LFP gamma for all cell types. Fast-spiking interneurons are moderately phase locked to LFP gamma for up to 3 spikes per theta stimulation period.

**Figure S2:**
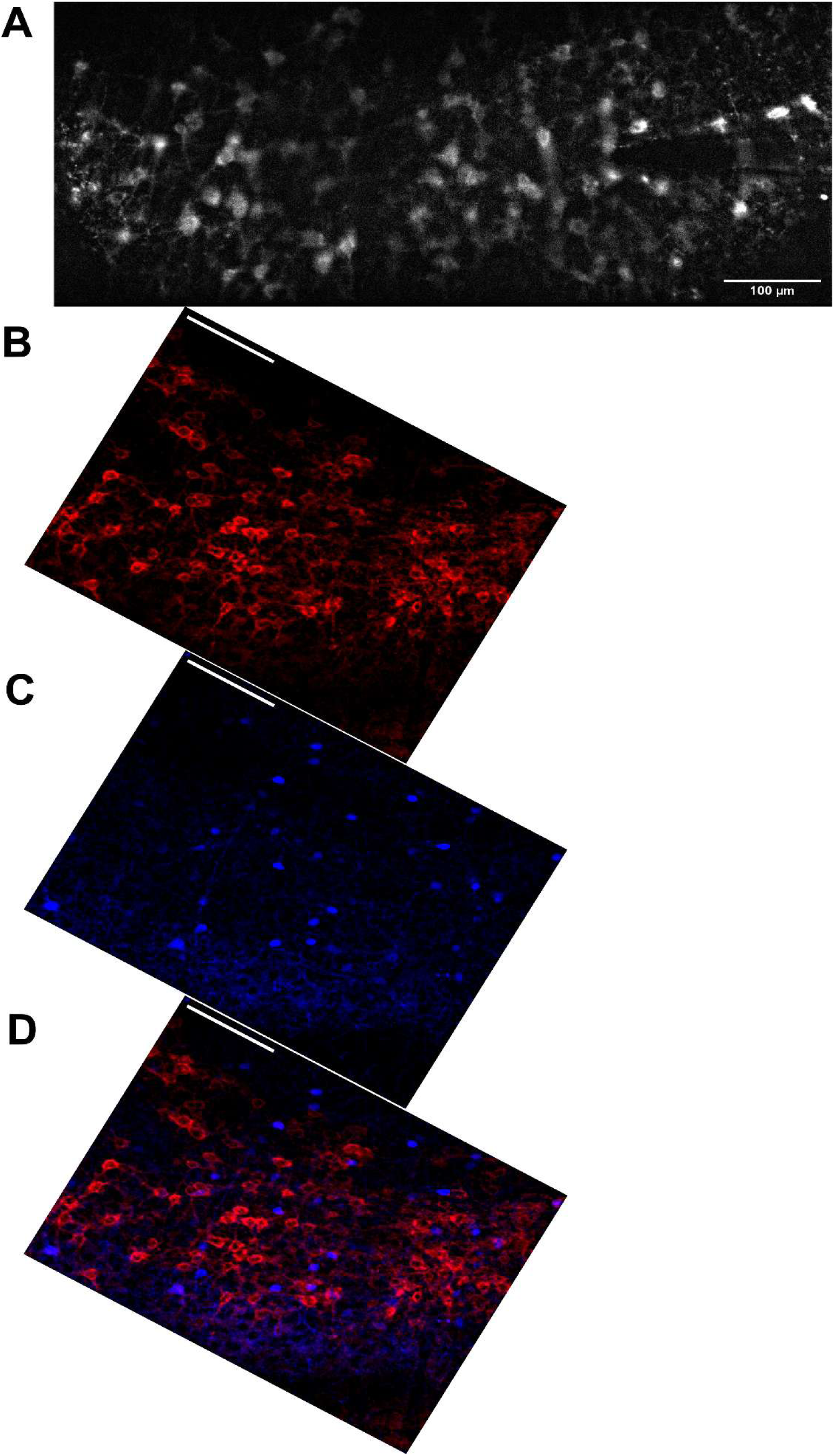
Minimal PV+ overlap with Voltron2 expression in layer II/III mEC. A) Example voltage imaging field of view with dense Voltron2 expression. B) Histological image of Voltron2-JF585 fluorescence registered to experimental field of view. C) Immunohistological image of PV expression in the same field of view as A and B. D) Composite image of Voltron2-JF585 (red) and PV (blue) expression in layer II/III mEC demonstrating minimal overlap between the populations.

**Figure S3:**
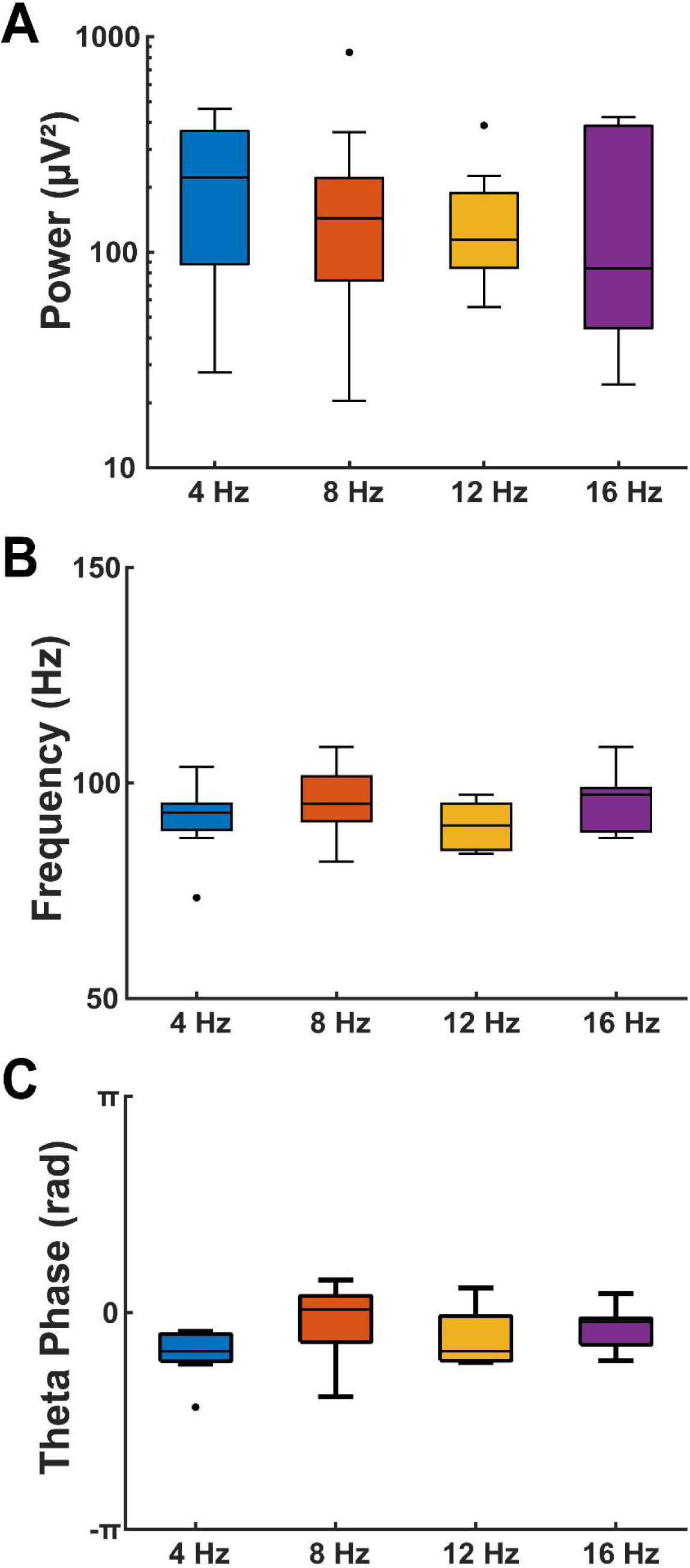
LFP gamma frequency activity doesn’t change with theta stimulation frequency. A) Average scalogram peak LFP gamma power from each voltage imaging field of view. B) Average scalogram LFP gamma frequency. C) Average scalogram LFP theta phase of peak gamma power.

**Figure S4:**
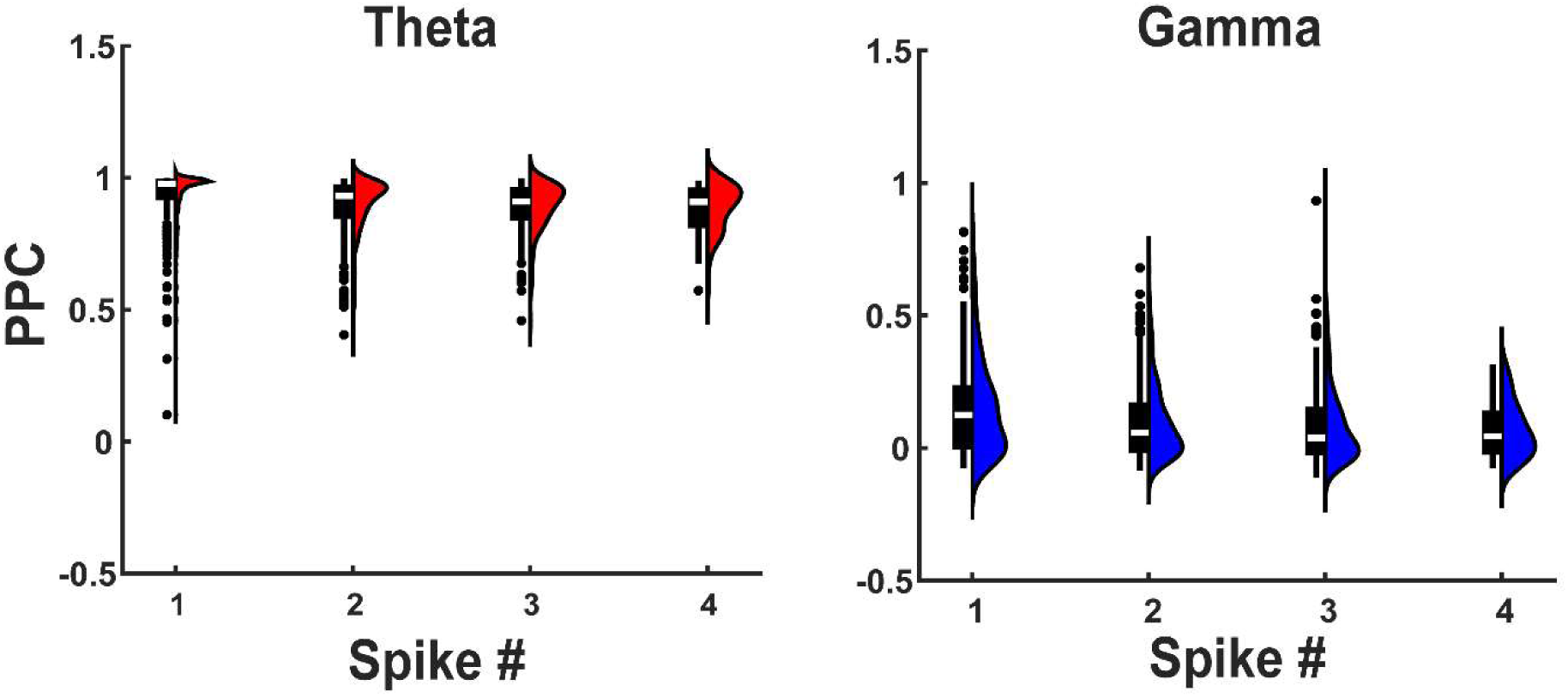
Strong theta (left), but not gamma (right) phase locking across multiple consecutive spikes per theta period from neurons in all imaging sessions.

**Figure S5:**
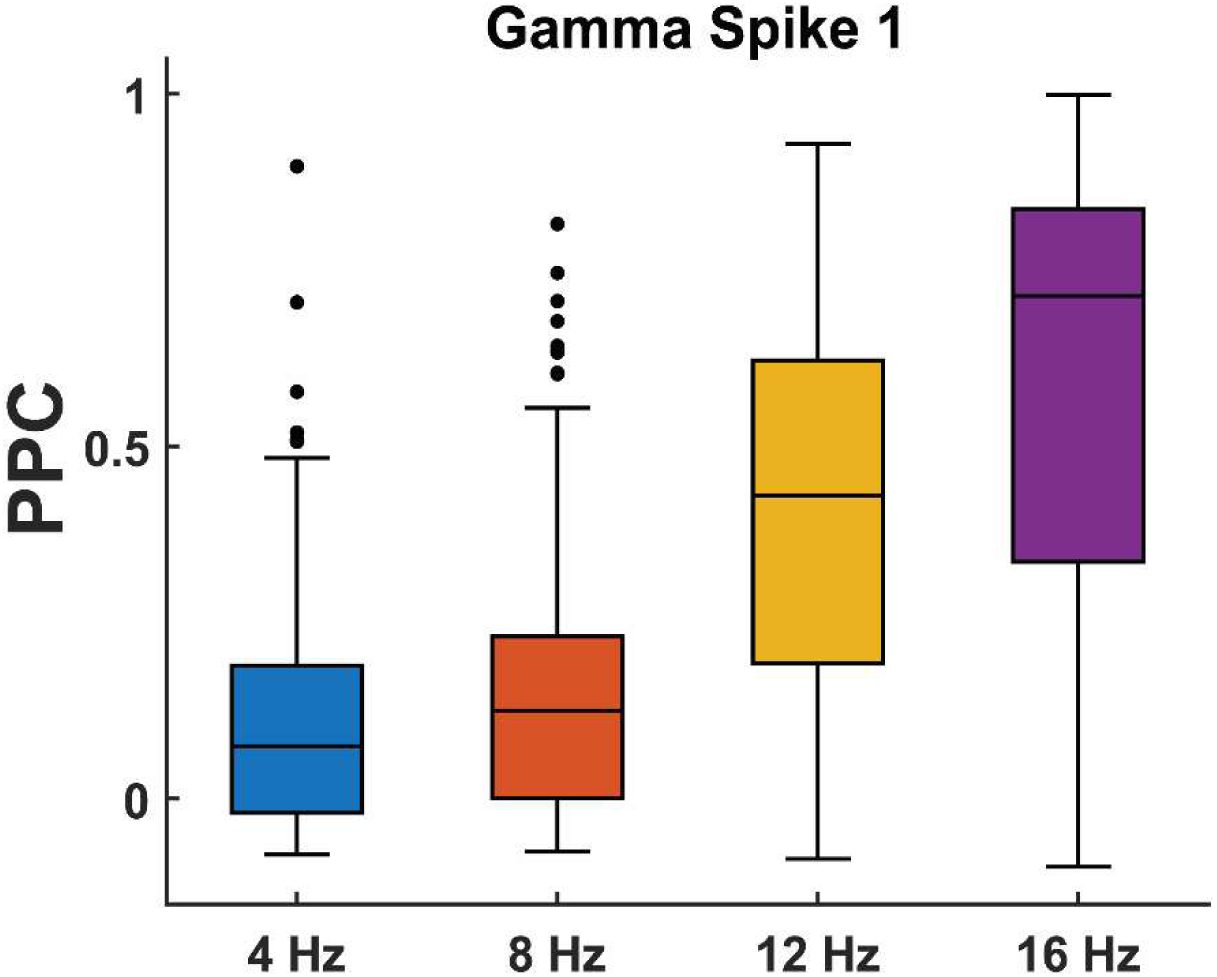
First spike gamma phase locking increases with theta stimulation frequency.

**Figure S6:**
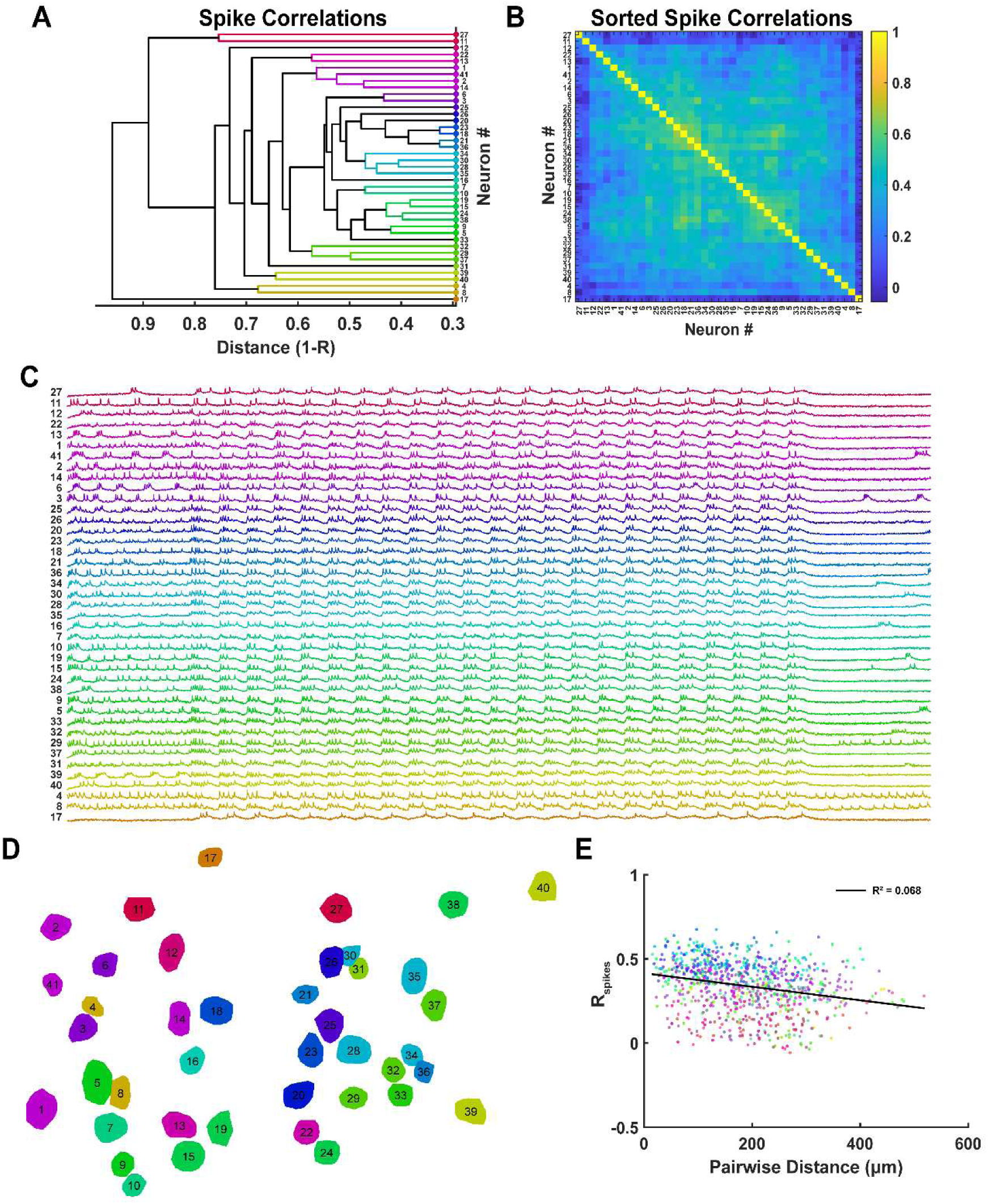
Spiking activity is not clustered across mEC. A) Agglomerative dendrogram of average spike correlation linkages between 41 mEC neurons in Fig. 5, 6. B) Sorted spike correlation matrix based on hierarchical clustering optimal leaf order. C) Sorted time series voltage activity based on hierarchical clustering of spike correlation matrix. D) Spatial organization of spike correlation clusters across the imaging field of view. E) Pairwise spike correlations are not correlated with distance.

